# Recombinant *Lactococcus lactis*-based multivalent vaccine targeting *Campylobacter jejuni* colonisation and modulating cecal microbiota in poultry

**DOI:** 10.1101/2025.09.27.678929

**Authors:** Prakash Biswas, Sakil Ahmed, Samiran Mondal, Samson Oladokun, Ozan Gundogdu, Amirul Islam Mallick

**Affiliations:** Department of Biological Sciences, Indian Institute of Science Education and Research Kolkata, Mohanpur, Nadia, 741246, West Bengal, India; Department of Veterinary Pathology, West Bengal University of Animal and Fishery Sciences, Kolkata, West Bengal 700037, India; Department of Poultry Science, Texas A&M University, College Station, TX 77843, USA; Department of Infection Biology, Faculty of Infectious & Tropical Diseases, London School of Hygiene and Tropical Medicine, London, WC1E 7HT, United Kingdom

**Author notes:** Corresponding Authors, Dr. Amirul Islam Mallick, DVM, PhD, Professor, Department of Biological Sciences, Indian Institute of Science Education and Research Kolkata, Mohanpur, Nadia, West Bengal, 741246, India. -Ext 1221, Dr. Ozan Gundogdu, PhD, Associate Professor, Department of Infection Biology, Faculty of Infectious & Tropical Diseases, London School of Hygiene and Tropical Medicine, London, WC1E 7HT, United Kingdom.

**Keywords:** *Campylobacter jejuni*, Chickens, *Lactococcus lactis*, Chitosan, Multivalent vaccine, Gut health

## Abstract

**Background:** Diarrhoeal diseases remain the second leading cause of preventable death globally, particularly among children under the age of five in developing countries, accounting for an estimated 2-3 million deaths annually. Among the bacterial pathogens causing diarrheal illness, *Campylobacter jejuni* (*C. jejuni*) remains one of the major contributors, particularly in LMICs. As a common gut pathogen, *C. jejuni* expresses several secretory or surface-expressed colonisation proteins, namely Hcp, VgrG, CadF, FlpA, and JlpA. Most of these proteins play pivotal roles in bacterial self-survival, host-cell adhesion, and invasion in avian and non-avian hosts.

**Methods:** To minimise the *C. jejuni* adhesion and subsequent colonisation in the avian gut, we explored the potential of a multivalent mucosal vaccine composition using these putative subunits of *C. jejuni*. For this purpose, we bioengineered a food-grade Lactic Acid-producing Bacterium, *Lactococcus lactis* (*L*. *lactis*), to express three key immunogenic subunits, Hcp, CadF and JlpA. Utilising this live vector-based multi-component mucosal vaccine platform, we investigated the immunoprotective potential of these antigens in chickens. Since the particular strain of *L. lactis* is non-colonising, we used chitosan as a natural mucoadhesive, biodegradable polymer to microencapsulate the engineered bacteria to increase their gut retention time for optimal interaction with local immune cells.

**Results:** Our *in vivo* immunisation study demonstrated that oral administration of this multivalent vaccine formulation elicited a strong local antibody response (sIgA) and upregulated key pro-inflammatory cytokines, leading to robust mucosal immune protection against the cecal colonisation of *C. jejuni*. Moreover, gut metagenomic analysis of vaccinated birds revealed a marked reduction in the phylum Campylobacterota, accompanied by an increased abundance of the phyla Bacillota and Bacteroidota, as part of a beneficial microbial community.

**Conclusions:** Together, this study underscores the potential of a live vector-based, multivalent mucosal vaccine as a promising, cost-effective strategy to reduce the risk of foodborne transmission of *C. jejuni*, particularly in poultry production systems.

## INTRODUCTION

Diarrhoeal diseases are the second leading cause of preventable death globally, particularly affecting children under the age of five in developing countries, with an estimated 2-3 million deaths annually [1–3]. Bacterial diarrhoeal illnesses are primarily transmitted via the faeco-oral route and are commonly caused by *Escherichia coli*, *Yersinia enterocolitica, Salmonella*, *Shigella*, *Vibrio cholerae*, and *Campylobacter jejuni* (*C. jejuni*) [4–8]. Among these pathogens, *C. jejuni* is recognised as the leading bacterial cause of human diarrhoeal disease and is especially prevalent in low- and middle-income countries (LMICs) [9–12]. Although *C. jejuni* infects a wide range of hosts, poultry, particularly chickens, are the primary reservoir and a significant source of human infection [13–15].

Recently, we demonstrated the role of key secreted and surface-expressed colonisation proteins (SECPs) of *Campylobacter jejuni*, such as Hcp and JlpA, in mediating host cell adhesion, cellular invasion, and subsequent pathogenesis in both avian and non-avian hosts [16,17]. Furthermore, mucosal administration of either Hcp or JlpA modestly reduced *C. jejuni* cecal colonisation in chicken and murine models [17,18]. Given that cecal colonisation by *C. jejuni* is a multifactorial process involving several putative virulence factors, we explored the efficacy of a multivalent vaccine formulation comprising JlpA (lipoprotein that binds explicitly to Hsp90α), Hcp (a key effector protein of bacterial T6SS), and CadF (a critical adhesion protein that facilitates host cell binding via fibronectin) [19–22]. Moreover, we hypothesised that since the sequences of the selected target genes (CadF, Hcp, and JlpA) are highly conserved across *C. jejuni* strains, if used in combination, they may ensure “broad-spectrum” immune protection against *C. jejuni* [23–25].

Building on this hypothesis, we employed our previously established mucosal vaccine delivery platform using a food-grade Lactic Acid-producing Bacterium (LAB) to surface express the CadF, Hcp, and JlpA proteins of the *C. jejuni* ^18^. The specific strain used for this study was *Lactococcus lactis* subsp. *cremoris* (MG1363), a GRAS (Generally Recognised As Safe) category LAB vector [27]. Given that this strain of *L. lactis* (also known as NZ9000) is non- colonising by nature, we utilised chitosan (CS), a natural, mucoadhesive, and biodegradable polymer, to microencapsulate the recombinant *L. lactis* (r*L. lactis*) cells [28,29]. This encapsulation aimed to enhance the stability and gut retention of *L. lactis* and promote optimal interaction with local immune cells, thereby improving the overall mucosal immunogenicity against the target protein [29]. Finally, the mixture of CS-coated r*L. lactis* in optimal numbers (3×10^9^ CFU) was orally administered (oral gavage) in seven-day-old *Campylobacteria*-free chicks (Rhode Island Red) for three consecutive weeks, followed by a challenge infection with a highly colonising strain of *C. jejuni* (TGH 9011) [30].

We demonstrated that oral administration of this multivalent vaccine formulation elicited a robust local immune response, characterised by a marked increase in secretory IgA (sIgA) levels and triggered the expression of key pro-inflammatory cytokines (such as IL-8, IL-1β, IL-17). Histopathological analysis of the challenged birds immunised with the present vaccine compositions further supported the positive impact of the combinatorial vaccine strategy using a LAB-based delivery platform on overall gut health. These findings align with the general health benefits of probiotics in inhibiting the growth of colonising pathogens by competitive adhesion on the epithelium and producing antimicrobial substances (like bacteriocins), especially by LAB vectors [31–33].

Since *Campylobacter jejuni* causes little or no pathogenicity in the chicken gut, we were interested in assessing how the bioengineered LAB vector, expressing heterologous proteins, influences the gut microbiome [34,35]. We employed full-length 16S rDNA gene sequencing (∼1500 bp) using long-read sequencing technology to achieve higher taxonomic resolution of microbial populations [36]. Our metagenomics data and the *in vivo* challenge experiment reveal an overall reduction in *Campylobacter* spp., accompanied by an increased abundance of *Lactobacillus* spp. and other beneficial microbiota, compared to unvaccinated control birds. Our metagenomic data and histopathological analysis indicate improved gut health and a more favourable microbial composition, which is at least partly attributable to the intrinsic probiotic effects of the LAB vector used as a vaccine delivery platform.

Together, the present multivalent vaccine against *C. jejuni* demonstrates significant benefits, including a marked reduction in *C. jejuni* colonisation, highlighting the effectiveness of targeting multiple virulence factors for enhanced protective immunity.

## MATERIAL AND METHODS

### Bacterial strains, cell lines, and other reagents

#### Bacteria

The list of bacterial strains and plasmids used in this study is presented in **Table S1 (Supporting Information)**. *Lactococcus lactis* subsp. *cremoris* NZ9000 cells and all recombinant *L. lactis* strains were routinely cultured at 30 °C in GM17 broth (HiMedia, India) without shaking. When required, chloramphenicol (SRL, India) was used with a final 20 µg/mL concentration for recombinant strains. The recombinant *E. coli* strains (TOP10), harbouring plasmids, were cultured in Luria-Bertani (LB) broth (Himedia) at 37 °C containing 20 µg/mL of chloramphenicol [18]. *C. jejuni* TGH 9011 was obtained through BEI Resources, NIAID, NIH: *Campylobacter jejuni* subsp. *jejuni*, Strain TGH 9011, NR-4082. *C. jejuni* was cultured at 37°C in Muller-Hinton (MH) broth (Himedia) supplemented with CAT selective supplement (Himedia) in a tri-gas incubator.

#### Cell lines

19-day-old embryonated eggs were used to isolate primary chicken embryo intestinal cells (CEICs) and cultured as described earlier [37]. For CEICs, intestinal sections of 19-day-old embryos were aseptically collected and cut into small pieces in 1×PBS. After that, 0.25% trypsin-EDTA was added, and the tissue was stirred for 45 min at room temperature (RT). Cells were isolated from the suspension through a cell strainer (40 µm) and centrifuged at 500g for 10 min at RT. Isolated primary CEICs were maintained in RPMI 1640 (Gibco, Invitrogen) medium supplemented with 10% FBS, 100 μg/mL streptomycin, and 100 U/mL penicillin at 37°C with 5% CO_2_ for 4-6 days until confluency reached 70-80%.

#### Chemicals and reagents

The reagents and chemicals used in this study were of the highest purity available. Chitosan was purchased from HiMedia Laboratory, and Sodium Tripolyphosphate (TPP) was procured from Loba Chemie (India).

### Generating recombinant *L. lactis* (r*L. lactis*) surface expressing CadF, Hcp and JlpA protein of *C. jejuni*

For r*L. lactis* surface expressing *C. jejuni* JlpA and Hcp antigen, we used our previously constructed vector with pNZ8048 plasmid backbone [17,18]. For the CadF (912 bp), the sequence without the signal peptide was amplified by PCR from the genomic DNA of *C. jejuni* (BCH71) **(Table S4, Supporting Information)**. The restriction-digested amplicon was cloned in between downstream of a signal peptide (_SP_) of *L. lactis* protease USP45 (_SP_-USP45; 81 bp, includes 27 residues USP45 leader peptide and a cleavage site for signal peptidase) and upstream of cell wall anchor motif of M6 gene from *Streptococcus pyogenes* (CWA_M6_; 424 bp) as per published method [17,18]. GenePulser (BioRad, USA) was then used to electrotransform the recombinant plasmid into *L. lactis* cells as per the method described previously [38]. Finally, the expression of the target protein was induced by the optimal concentration of Nisin (Sigma, USA) (15 ng/mL)[39] **(Figure S1, Supporting Information)**.

### Chitosan (CS) coating of *L. lactis* and r*L. lactis* surface expressing CadF, Hcp and JlpA protein

Approximately 4 h post-induction with nisin, *L. lactis* and r*L. lactis* cells (∼3 × 10[CFU) were centrifuged at 2400g for 5 min to collect the bacterial pellet. Then, the bacterial pellet was dissolved in 1× PBS (700 µL), added to 0.05% (w/v) CS solution (3.3 mL), and stirred for 30 min at RT. After incubation, 1 mL of 0.1% (w/v) sodium tripolyphosphate (STPP) solution was added dropwise and further incubated for 1 h under stirring conditions at RT. Then, the solution was pelleted and washed thrice with 1× PBS to remove residual unpolymerised CS and STPP using centrifugation at 2400g for 5 min. Finally, the coated r*L. lactis* cells were dissolved in 1× PBS for further assessment [37].

#### Determining the accessibility of the recombinant protein expressed by CS-coated rL. lactis

To check the accessibility of surface-expressed protein, 3×10^9^ CS-coated r*L. lactis* cells were prepared for antigen-specific primary antibody coating. For this, the cells were first blocked with 3% BSA and fixed with pre-chilled 4% PFA. The cells were then washed and probed with anti-CadF/Hcp/JlpA antibodies raised in rabbit (1:100 dilution) overnight at 4°C. After incubation, cells were washed with PBS and incubated with FITC-conjugated goat anti-rabbit IgG antibody (H + L) (1:500 dilution) (Life Technologies) for 2 h at room temperature. Finally, the cells were washed and subjected to flow cytometric analysis in BD LSR Fortessa using the FITC filter (Excitation/Emission at 488/535 nm) [40,18,37].

*In vitro cell adhesion of CS-coated* r*L. lactis*

The mucoadhesive property of chitosan-coated r*L. lactis* expressing rCadF, rHcp, and rJlpA proteins to primary chicken embryo intestinal cells (CEICs) was assessed using confocal microscopy. Briefly, nisin (15 ng/mL) induced uncoated or CS-coated r*L. lactis* cells were incubated with CEICs at a ratio of 1:1000 (CEICs: bacteria) for 4 h at 37°C. Then, cells were washed with PBS and fixed with 4% PFA. After 15 min, cells were thoroughly washed and blocked using 3% BSA for 1 h at RT, followed by probing with an antigen-specific primary antibody raised in rabbit (1:100 dilution) for 2 h at RT. After three washes, cells were stained with FITC-conjugated goat anti-rabbit IgG (H + L) secondary antibody (1:500 dilution) (Thermo Fisher Scientific, USA). Further, the cells were stained with phalloidin 647 and DAPI and mounted onto a glass slide with Vectashield mounting media (Vector Laboratories, USA) for confocal microscopy and images were captured (405 nm laser for DAPI; 488 nm laser for FITC; 638 nm laser for Phalloidin 647) [17,18,37,39,41,42].

### *In vivo* immunogenicity of mucosally administered r*L. lactis* in chickens

#### Immunogen preparation and experimental groups

Recombinant *L. lactis* displaying rCadF, rHcp, and rJlpA proteins were prepared using the optimal nisin concentration (15 ng/mL). To coat with chitosan-TPP, the induced cells were pelleted, washed with sterile endotoxin-free PBS, and resuspended in PBS. To assess the *in vivo* immunogenicity of mucosally (intragastric) administered *L. lactis* cells expressing rCadF, rHcp, and rJlpA proteins, a total of 120 Rhode Island Red (RIR) chicks were purchased from ARD RKVY Haringhata Farm (India). The experimental birds were maintained in a deep litter system throughout the trial and fed an *ad libitum* antibiotic-free corn-soybean-based mash diet. The detailed composition of the substrate used in this study is provided in **Table S2 (Supporting Information)**. On day 7, the chicks were randomly divided into seven experimental groups, with 24 birds each. Chicks belonging to different groups received the following treatments in 100µL of PBS. Control Group (PBS): 100µL PBS/bird; Group CS: 100µL CS-TPP/bird; Group *L. lactis*: 3×10^9^ CFU *L. lactis* NZ9000/bird; Group r*L. lactis* (CadF): 1×10^9^ CFU r*L. lactis*-CadF with 2×10^9^ CFU NZ9000/bird; Group r*L. lactis* (JlpA): 1×10^9^ CFU r*L. lactis*-JlpA with 2×10^9^ CFU NZ9000/bird; Group r*L. lactis* (Hcp): 1×10^9^ CFU r*L. lactis*-Hcp with 2×10^9^ CFU NZ9000/bird; Group r*L. lactis* (CadF+JlpA+Hcp): 1×10^9^ CFU r*L. lactis*-CadF+ 1×10^9^ CFU r*L. lactis*-JlpA+ 1×10^9^ CFU r*L. lactis*-Hcp/bird.

#### Sample collection

At day 7 post-last feeding (day 28), half of the birds from each group were sacrificed to collect the Bursa of Fabricius (BOF), cecal tonsils, spleen, blood, faeces, and intestinal lavages. In contrast, the remaining (n = 12) birds from each group were challenged with 1×10^8^ CFU of *C. jejuni* TGH 9011 (ATCC 43431). All challenged birds were euthanised on the seventh post-challenge day (day 35). Birds were euthanised by CO_2_ asphyxiation (30%) followed by cervical dislocation as per the guidelines approaved by Committee for the Purpose of Control And Supervision of Experiments on Animals (CCSEA), Ministry of Fisheries, Animal Husbandry and Dairying Department of Animal Husbandry and Dairying, Govt. of India (Source: Manual for CCSEA Guidelines For Polutry/Birds Facility, 2020).

For every bird in each group, samples of the BOF and spleen weighing around 100 mg were collected and stored in Trizol at -80°C. Throughout the experiment, weekly samples of blood and faeces were collected from individual birds in each group. Faecal pellets were processed using the previously mentioned procedure, while serum samples were aliquoted and kept at - 20°C until further use [43]. Intestinal lavages were harvested following a published protocol standardised in our lab and stored at -20°C [18].

### Assessing local antibody responses in immunised chickens

**Secretory IgA (sIgA) titer in intestinal lavages and faecal pellets:** To evaluate the sIgA level against whole cell lysate (WCL) of *C. jejuni* TGH 9011 in clarified gastric lavage or faecal samples, indirect ELISA was performed using half-diluted samples (as starting dilution). Briefly, 1 µg/well of *C. jejuni* WCL was used to coat the ELISA plates overnight at 4°C. The next day, plates were washed with PBST (0.1% Tween 20) and blocked using 5% BSA for 1 h. After thoroughly washing, intestinal lavages or faecal soup samples were added and incubated at RT for 2 h. Plates were then washed using PBST and probed with goat anti-chicken IgA HRP-conjugated as the secondary antibody (1: 3000 dilutions; Bethyl Laboratories) at RT for 1 h. Following washes with PBST, 3,3’,5,5’-Tetramethylbenzidine (TMB) substrate was added to each well, and the reaction was stopped with 1 M H_2_SO_4_ after 5 min of incubation. Finally, the absorbance was read using a microplate reader (BioTek) at 450 nm [18].

### Assessment of cellular responses in immunised birds

#### Isolation of splenocytes

The spleens of five experimental birds (n=5) per group were collected, and a single-cell suspension was prepared following the method described in previous studies [17,44]. In brief, the spleens were minced in a sterile petri dish containing RPMI 1640 (Gibco, USA) medium using a disposable syringe plunger. The cell suspension was aspirated and passed through a 40 µm cell strainer (Corning, Merck). The resultant filtrate, comprising the single-cell suspension, was placed onto pre-warmed Histopaque-1077 solution (Sigma) in a 1:1 ratio and centrifuged at 500g for 20 min at room temperature. The splenocyte interface was carefully harvested, washed, and resuspended in a complete growth medium (RPMI 1640).

### *In vitro* Nitric Oxide (NO) production

To assess the level of nitric oxide (NO) production by antigen-primed splenocytes from experimental birds, a standard Griess assay was conducted according to the manufacturer’s instructions (Sigma). Briefly, a single-cell suspension of splenocytes was seeded in 24-well tissue culture plates for three hours at a density of 1.5× 10^5^ in phenol red-free complete RPMI 1640 growth media. Then, splenocytes were challenged with *C. jejuni* WCL (1 μg/mL). Following 48 h of incubation, 100 μL of culture supernatant was collected from each well and incubated with an equal volume of Griess reagent at room temperature for 15 min. The absorbance for each well was measured at 540 nm using a microplate reader (Biotek, Epoch2) [45,46]. The concentration of nitrite was determined in comparison with a standard curve generated using sodium nitrite (NaNO_2_) (Sigma) **(Figure S2, Supporting Information)**.

### *In vitro* splenocyte proliferation

The splenocyte proliferation assay was then conducted using Click-iT Plus EdU Alexa Fluor 488 Flow Cytometry Assay Kit (Invitrogen) according to the manufacturer’s protocol. Briefly, 3×10^5^ cells in RPMI 1640 medium were seeded into each well of a 12-well tissue culture plate. After 3 h, the cells were stimulated with 10 µg/mL of *C. jejuni* TGH 9011 whole-cell lysate (WCL). Unstimulated splenocytes and splenocytes stimulated with 10 µg/mL of concanavalin A (Con A) were used as controls. After 24 h of stimulation, 10 µM of 5-Ethynyl-2′-deoxyuridine (EDU) was added to the wells and incubated for an additional 24 h. The cells were then washed with PBS containing 0.2% BSA and fixed for 15 min on ice. Following fixation, the cells were rewashed and permeabilised for 15 min on ice. The cells were subsequently incubated on ice with EDU buffer additive and reaction cocktail for 30 min. After incubation, the cells were washed and analysed by BD LSRFortessa flow cytometer (BD Biosciences) [47]. The splenic lymphocyte proliferation index (Stimulation Index; SI) for each experimental group was calculated using the following formula: Stimulation Index (SI) = Mean fluorescence of stimulated cells / Mean fluorescence of unstimulated cells [48].

### Analysis of B and T cell subsets in cecal tonsils and Bursa of Fabricius (BOF)

#### Isolation of mononuclear cells from intestinal tissue

Whole cecal tonsils and a portion of the Bursa of Fabricius (BOF) were collected from chickens and kept on ice in PBS containing penicillin (10 U/mL) and streptomycin (10 µg/mL). Each tissue was then sectioned into smaller pieces and washed three times with PBS. Tissue samples were then digested with collagenase type I (4 mg/mL at 37°C for 20 min; Merck) in PBS containing penicillin (100 U/mL) and streptomycin (100 µg/mL). The resulting tissue digests were filtered through 40-µm cell strainers (Corning, Merck), using the rubber end of a 10 mL syringe plunger to crush the tissue. The cell suspensions from the cecal tonsils and BOF were layered onto Histopaque-1077 (Sigma) in a 1:1 ratio, followed by density-gradient centrifugation at 500g for 20 min to isolate the mononuclear cells. The aspirated buffy coats were washed by centrifugation at 400g for 5 min in RPMI 1640 medium containing penicillin and streptomycin. The mononuclear cells were then resuspended in a complete RPMI cell culture medium, which consisted of RPMI 1640, 10% FBS (Gibco), penicillin (100 U/mL), and streptomycin (100 µg/mL). Cell count and viability were assessed using a haemocytometer and the trypan blue exclusion method. The mononuclear cells were resuspended in a complete RPMI medium at a 1×10^6^ cells/mL density and kept on ice [28].

#### Flow cytometric analysis of B and T cell subsets

Mononuclear cells from cecal tonsils and BOF (1×10^6^ cells) were resuspended in FACS staining buffer (PBS containing 1% BSA) and subjected to staining with specific antibody panels. Cells were labelled with mouse anti-Bu1- FITC and mouse anti-IgA-PE to characterise B cells. For T cell identification, the cells were stained with mouse anti-CD3ζ-APC, mouse anti-CD4-FITC, and mouse anti-γδTCR-PE. The cells were incubated with the respective antibodies for 15 min at 4°C in FACS staining buffer. After the staining procedure, cells were washed at 400g for 5 min with FACS staining buffer and then incubated for an additional 10 min at 4°C with 7-AAD (Southern Biotech, Canada). Following the incubation, cells were fixed with 4% paraformaldehyde (PFA) at 4°C. After fixation, cells were washed and stored at 4°C in FACS staining buffer [28]. Data was acquired on a BD LSRFortessa flow cytometer (BD Biosciences), collecting 5×10^4^ events. Flow cytometry data were subsequently analysed using FlowJo V10 software. A detailed list of the antibodies used is provided in **Table S3 (Supporting Information)**.

### Assessing cytokine gene expression in spleen and BOF tissues by qRT-PCR

According to the manufacturer’s instructions, total RNA was isolated from 50 mg of spleen and BOF tissues using Trizol reagent (Invitrogen, USA). cDNA synthesis for each sample was carried out using the iScript cDNA synthesis kit (Bio-Rad). To assess the expression of cytokine genes and transcription factors, qRT-PCR was performed using specific primers for chicken IFN-γ, IL-8, IL-1β, IL-17A, TNF-α, NF-κB and iNOS with the chicken β-actin gene serving as the internal control. Primer details are provided in **Table S4 (Supporting Information)**.

### *In vitro* functionality of sIgA against *C. jejuni* adherence to host cells

To evaluate the ability of intestinal sIgA to inhibit *C. jejuni* adherence and invasion, an *in vitro* protection assay was performed [44]. Briefly, 1.5×10^4^ *C. jejuni* cells were incubated with raw intestinal lavages for 3 h at 37°C. After this incubation, chicken CEICs were incubated with the treated *C. jejuni* at an MOI of 1:100 for 3 hours under 5% CO_2_. After incubation, the cells were washed and lysed with 1% Triton X-100. The lysates were serially diluted and plated onto *Campylobacter* selective agar plates, which were then incubated overnight at 37°C under microaerobic conditions in a tri-gas incubator (Thermo Fisher Scientific). Colonies that formed on the plates were counted for each experimental group.

### *In vivo* efficacy of mucosally administered r*L. lactis* against cecal colonisation of *C. jejuni*

To evaluate the protective efficacy of r*L. lactis*, birds from different experimental groups were challenged orally with 1×10^8^ CFU of *C. jejuni* TGH 9011 (in 100µL PBS) via oral gavage on day 7 following the last feeding. Seven days post-challenge, all birds were euthanised, and each bird’s cecal tissue and contents were collected.

#### Cecal load of C. jejuni

To determine the cecal load of *C. jejuni*, approximately 200 mg of cecal content from each bird was collected and dissolved in MH broth. Then, cecal contents were serially diluted in MH broth, plated onto *Campylobacter* selective agar supplemented with cefoperazone, amphotericin B, and teicoplanin (CAT) (Himedia), and incubated at 37°C in a tri-gas incubator. The number of bacterial colonies on the plate was counted using a colony counter (Scan 500, Interscience, France) and plotted for each group.

#### Histopathological changes in cecal tissues

On day 7 post-infection, cecal tissue from experimental birds challenged with *C. jejuni* was collected and processed for histological analysis in accordance with previously published protocols [49,50]. Briefly, 0.5 cm sections of cecal tissue were initially fixed in 10% formalin, followed by washing under running tap water. The tissue was then dehydrated through a graded series of acetone (70%, 90%, and 100%). After dehydration, the tissues were cleared with two changes of benzene and subsequently embedded in molten paraffin (62°C) through three 1 h changes. Following the sectioning of paraffin blocks, hematoxylin and eosin (H&E) staining was performed to make the slides.

### Effect of oral administration of r*L. lactis* in the gut microbial population

#### Full-length 16S rDNA sequencing of the cecal microbial community

Total genomic DNA was extracted from 250 mg of caecal content from each bird with the PowerFecal Pro DNA Kit (QIAGEN, Germany). For microbial community analysis, 30 ng high-quality genomic DNA was used as an input for the Nanopore 16S Barcoding Kit 24 V14 (SQK-16S114.24), which amplified the 16S rDNA (V1-V9 regions) gene by PCR using specific barcodes and LongAmp Hot Start Taq 2X Master Mix (NEB, M0533) for each sample following the manufacturer’s instructions. A total of 20 samples were sequenced (4 samples/group), and the amplicons were quantified using Qubit dsDNA HS Assay Kit (Invitrogen, Q32851), pooled in equimolar concentration and purified with AMPure XP beads (Beckman Coulter, USA). 50 fmol of the barcoded library was ligated with Nanopore adaptors supplied in the 16s kit and sequenced on the FLO-MIN114 flow cell using the MinION Mk1B sequencer (Oxford Nanopore Technologies, UK) [36,51].

#### Bioinformatics analysis of metagenomic data

The FASTQ files from Nanopore sequencing were base-called simultaneously with sequencing using Fast model v4.3.0, 400 bps with a minimum Q score of 8, with MinKNOW (Oxford Nanopore Technologies, UK). The barcoded samples were analysed using the wf-metagenomics workflow with EPI2ME v5.2.5 (Oxford Nanopore Technologies, UK). The sequences were referenced with the SILVA_138_1 database in the workflow. Then, a report of operational taxonomic units (OTUs) across all samples was generated. Further, the MicrobiomeAnalyst was used to analyse the relative abundance, alpha and beta diversity. The correlation network analysis of bacteria within the gut and pharyngeal microbiota was carried out by the SparCC method (r[<[0.3, P[<[0.05) using the plugin in MicrobiomeAnalyst [52,53].

### Statistical analysis

Graphical presentations and data analysis were performed using GraphPad Prism statistical software (Version 8.0.1). The Shapiro-Wilk test was applied to assess the normality of the data.

The Student’s t-test (two-tailed, unpaired) or the non-parametric Mann-Whitney U test was used to compare differences between experimental groups, depending on data distribution. Statistical significance was defined as *p ≤ 0.05, **p ≤ 0.01 and ***p≤ 0.001.

## RESULTS

### Nisin-induced uncoated or CS-coated r*L. lactis* showed stable surface expression of rCadF, Hcp and JlpA protein

To assess the accessibility of surface-expressed rCadF, rHcp, and rJlpA upon coating them with chitosan, induced r*L. lactis* cells were probed with polyclonal primary antibodies, followed by a FITC-conjugated goat anti-mouse/rabbit IgG (H+L) secondary antibody, and analysed by flow cytometry. A distinct shift in fluorescence intensity was observed in the gated population of uncoated and CS-coated induced r*L. lactis* cells for all three recombinant proteins compared to WT *L. lactis* NZ9000 (empty vector) cells **(Figure 1A, B and C)**.

**Figure 1:**
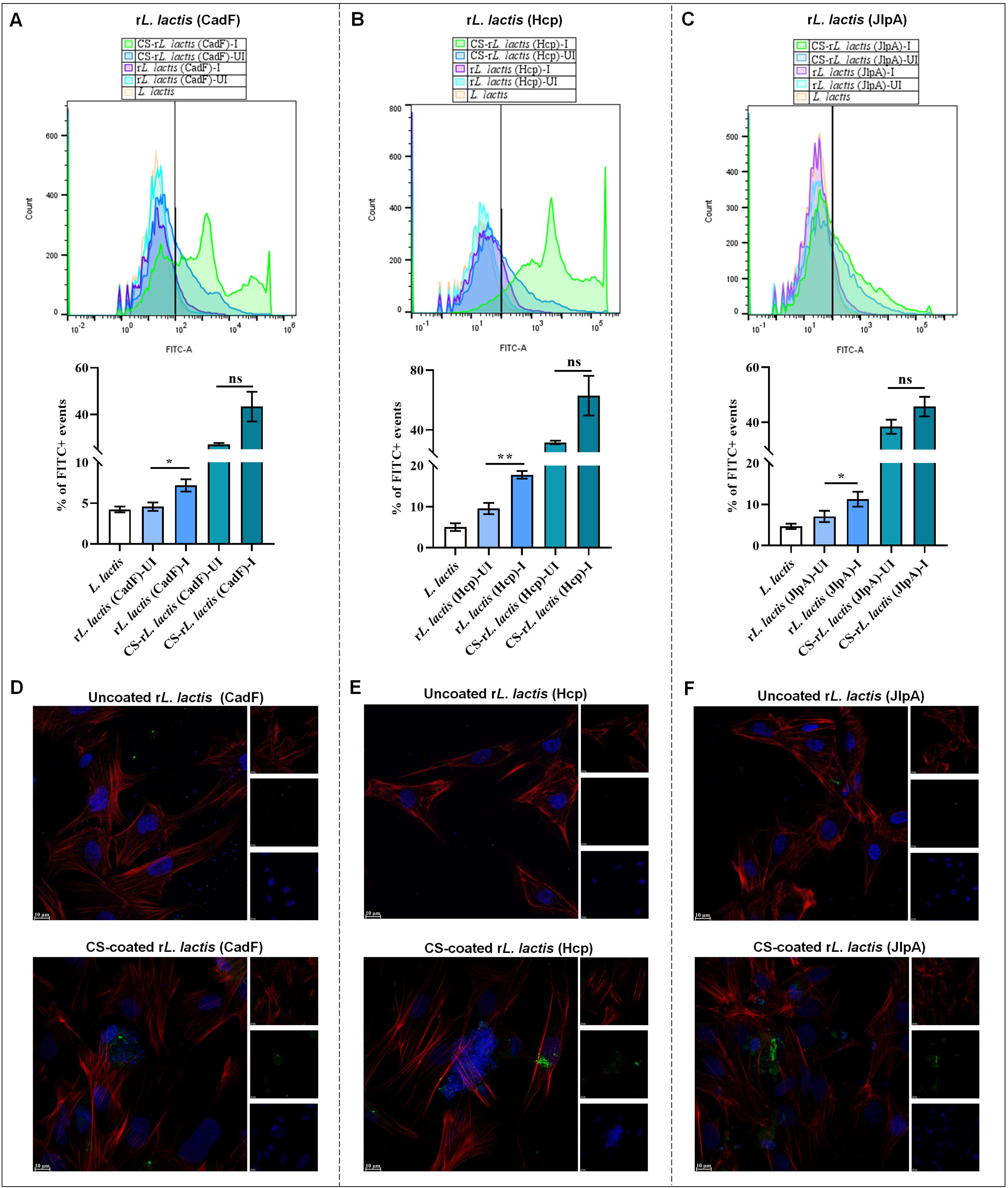
Nisin-induced CS-coated r*L. lactis* (CadF, Hcp and JlpA) surface expression and adhesion to host cells. **A-C :** Surface expression of CS-coated r*L. lactis* (CadF, Hcp and JlpA) by flow cytometry. Histograms denote the FITC-A signal vs counts in each group, and the bar graph for % of FITC+ events among the groups. Data shows enhanced signals in the CS-coated and uncoated induced (I) groups compared to the CS-coated and uncoated uninduced (UI) groups. Each bar represents the mean % of FITC+ events ± SE from two independent experiments (n=5). Asterisks indicate a statistically significant difference (**p ≤ 0.01) compared to the control group. **D-F:** *In vitro* adhesion assay of uncoated and CS-coated r*L. Lactis* (CadF, Hcp, JlpA) in primary chicken embryo intestinal cells (CEICs). The cells were incubated with induced uncoated and CS-coated r*L. lactis* 4 h at 37°C with 5% CO_2_, at a ratio of cell vs bacteria 1:1000. Representative CLSM images of cells incubated with CS-coated r*L. lactis* cells showed enhanced cell adhesion compared to uncoated bacteria. Scale bar: 10 μm.

### The CS-coating of r*L. lactis* enhances bacterial adhesion to host cells

We assessed the CS-coated r*L. lactis* cell adhesive performance to investigate whether electrostatic interactions between positively charged CS-coated bacteria enhance interactions with host cells. Confocal microscopy images showed that CS-coated induced r*L. lactis* expressing CadF/Hcp/JlpA had higher fluorescence signals and adhesive properties to primary CEICs than uncoated induced bacterial cells **(Figure 1D, E, and F)** and **(Figure S3, Supporting Information).**

### *Oral administration of CS-coated* r*L. lactis* induces significant local immune responses in chickens

#### Induction of local IgA (sIgA) responses

To evaluate the ability of mucosal delivery of r*L. lactis* expressing CadF, JlpA, and Hcp to induce an antigen-specific local antibody response, intestinal lavages and faecal samples were collected on day 7 post-last immunisation and analysed for sIgA **(Figure 2A)**. Following intra-gastric administration of CS-coated r*L. lactis* cells over three weeks (3 consecutive days/ week), a significant increase was observed in local antibody level (secretory IgA) in the intestinal lavage and faecal soup of the birds that received the combination of r*L. lactis* expressing all three proteins compared to the individual feeding regimens **(Figure 2B)**. However, no significant changes in serum IgY response were observed among the groups (data not shown) [18].

**Figure 2:**
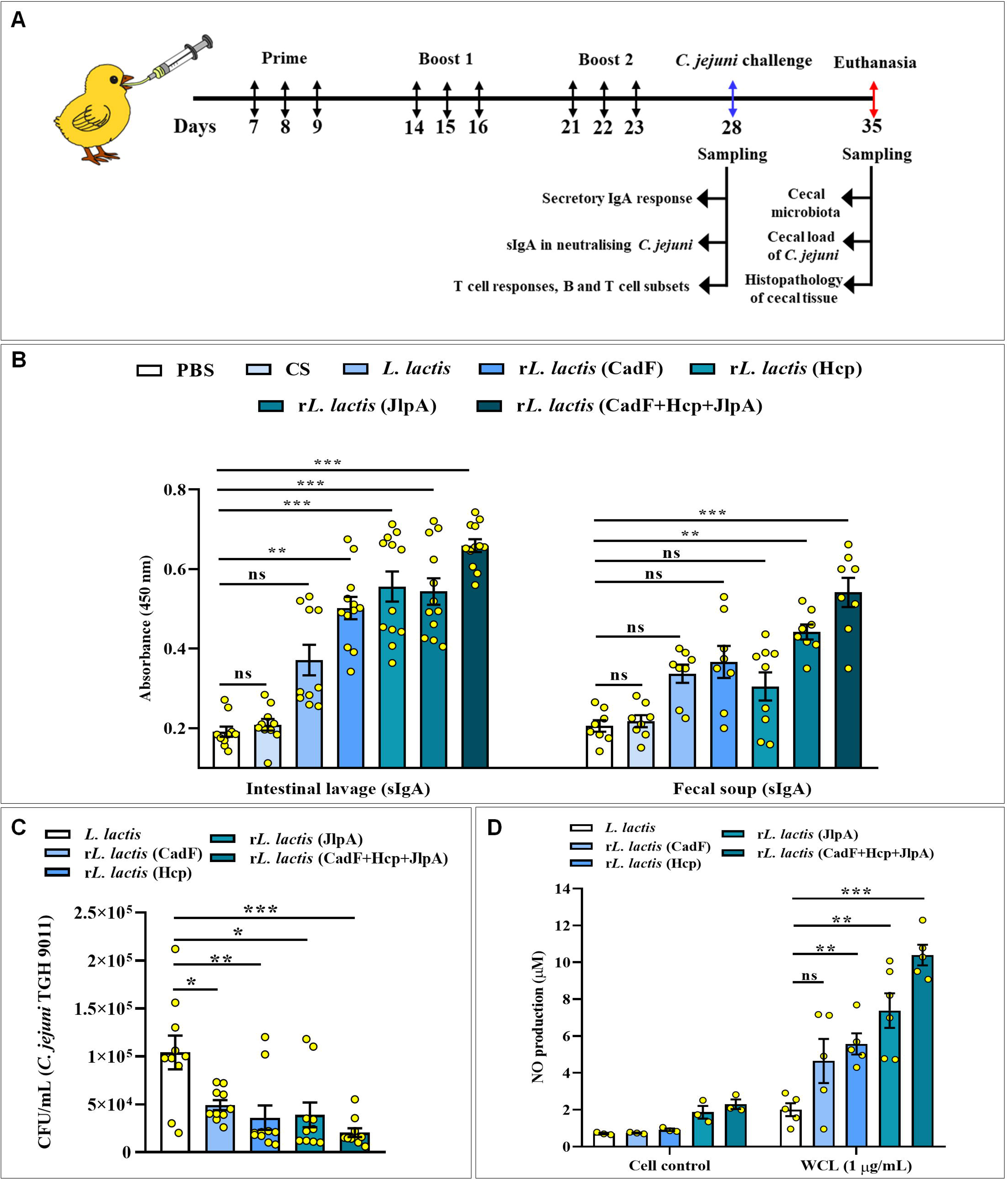
Schedule for oral administration of r*L. lactis* and assessment of antibody responses. **A:** Schematic of the chicken immunisation and sampling. 120-day-old chickens were orally administered (oral gavage) with r*L. lactis* as per the indicated time points for three weeks for two independent experiments (n=2). At day 7 post-first feeding (day 28), half of the birds were sacrificed to collect intestinal lavages, faeces, blood and tissue samples, while the remaining birds were challenged with 1×10^8^ CFU of *C. jejuni* (day 28). Challenged birds were sacrificed on day 35 post-first administration to collect cecal tissue and its content. **B:** Comparative analysis of mucosal (sIgA) antibody level in intestinal lavage, faecal soups collected from each bird belonging to different treatment groups. Data shows a significant increment of sIgA level in lavage and faecal soup of the birds administered with r*L. lactis* expressing CadF+Hcp+JlpA compared to the control group of birds (received PBS). Each bar represents the mean absorbance (A450) ± SE of 12 birds from two independent experiments. Asterisks indicate a statistically significant difference (***p ≤ 0.001) compared to the control group (PBS). **C:** *In vitro* neutralisation of *C. jejuni* adhesion and invasion of chicken primary CEICs by secretory sIgA present in the lavage sample of birds belonging to the different experimental groups, showing marked reduction in *C. jejuni* adhesion and invasion exerted by r*L. lactis* (CadF+Hcp+JlpA) administered group of birds (**p ≤ 0.01; immunised vs. control). Data represent mean CFU/mL ± SE (n = 10) from two independent experiments. **D:** High-level Nitric oxide (NO) production by the splenocytes collected from the birds administered with r*L. lactis* (CadF+Hcp+JlpA) when primed with whole cell lysate of *C. jejuni*. Data represent mean NO production ± SE (n =6).

### sIgA-mediated *in vitro* neutralisation of *C. jejuni* adhesion

A neutralisation assay was performed to assess the *in vitro* inhibitory effect of intestinal secretory IgA (sIgA) on *C. jejuni* adherence and invasion in CEICs. Direct comparison of total CFU of *C. jejuni* present in infected chicken embryo intestinal cells (CEICs) suggests that, compared to the individual antigen and controls, the sIgA present in the birds immunised with all three antigens significantly inhibits the host cell adhesion and invasion of *C. jejuni* (immunised vs. control) (p≤ 0.0079) **(Figure 2C)**.

### *In vitro* stimulation of splenic lymphocytes triggers cell proliferation and NO production

In response to *in vitro* stimulation with WCL, splenocytes from birds immunised with multi-component proteins (CadF, JlpA, and Hcp) exhibited significantly higher nitric oxide production in their culture supernatants compared to those immunised with individual proteins or empty NZ9000 cells (immunised vs. control) (p≤ 0.001). In contrast, birds treated with empty NZ9000 cells showed only a basal level of NO production **(Figure 2D)**.

The proliferation of specific splenocytes was significantly higher in birds immunised with the combination of all three antigens than in those receiving individual antigens or the control group (NZ9000), according to the splenocyte stimulation index **(Figure 3A)**.

**Figure 3:**
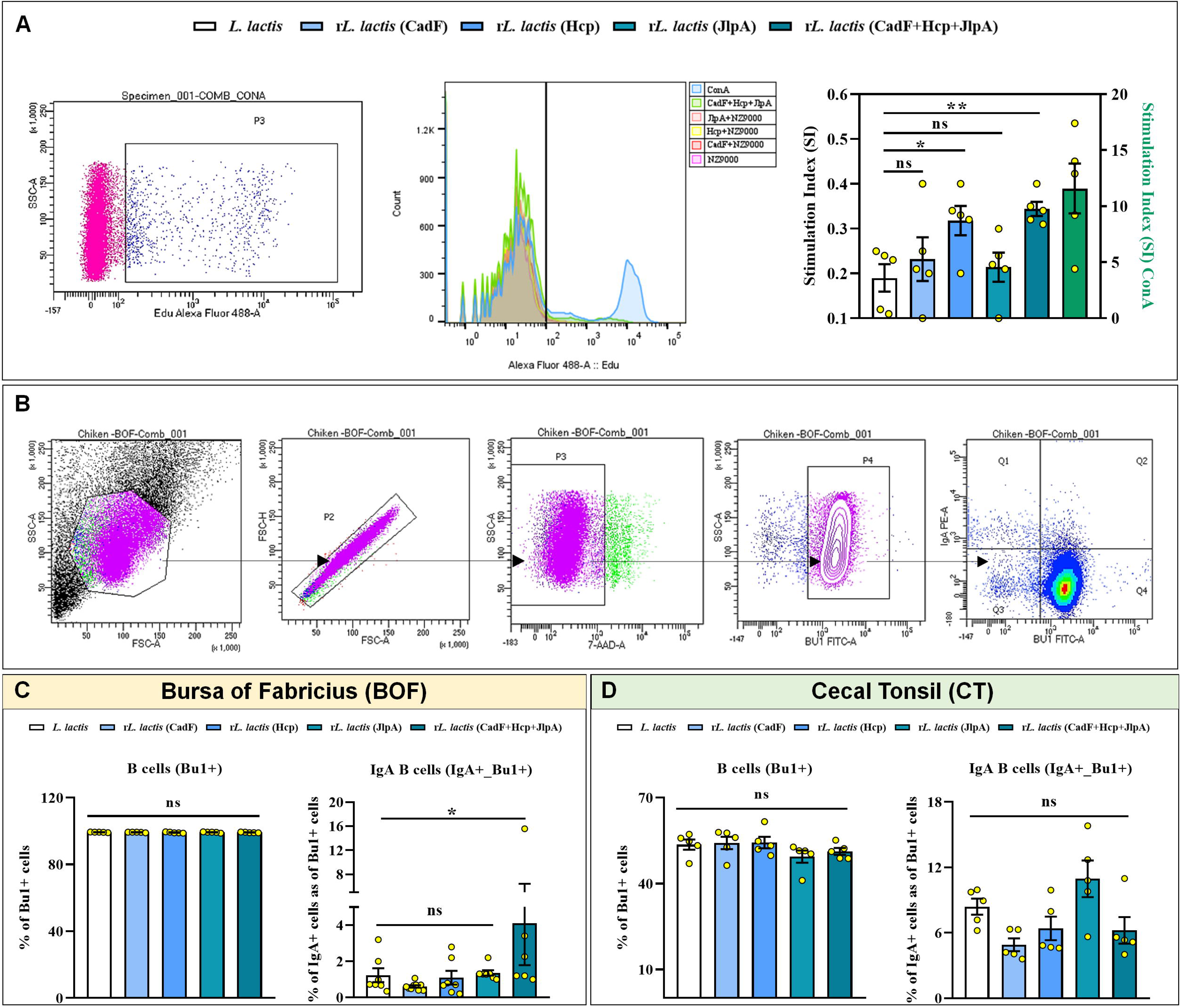
*In vitro* splenocyte proliferation *in situ* EDU assay and flow cytometry analysis of B cell population in BOF and cecal tonsils. **A:** *In vitro* splenocyte (lymphocyte) proliferation by *C. jejuni* whole cell lysate (WCL) (1 μg/mL of WCL) was measured by EDU assay performed at day 28 post first administration, showing a significant difference among r*L. lactis* (Cadf +Hcp+JlpA) compared to control groups (WT *L. lactis* only). Data represent mean stimulation index ± SE (n=5). Asterisks indicate a statistically significant difference (**p ≤ 0.01) from control (WT *L. lactis*). **B:** Panel showing gating strategy for IgA-positive B cells (Bu1+IgA+) present in the Bursa of Fabricius (BOF) and cecal tonsil (CT) of experimental birds. **C, D:** Flow cytometric analysis of IgA-positive B cells (Bu1+IgA+) suggests a modest increase in the birds administered with r*L. lactis* (CadF, JlpA and Hcp) compared to other experimental groups; however, no such changes could be observed in B cells present in CT.

### Change in the B and T cell phenotypes

Mononuclear cells were isolated from BOF, stained and analysed using flow cytometry following the gating strategy provided in **Figure 3B**. Compared to the control groups, a moderate increase in the total IgA-producing B-cells (Bu1^+^IgA^+^) was observed in the BOF of chickens that received r*L. lactis* expressing the combination of CadF, JlpA, and Hcp **(Figure 3C)**. However, no significant differences were observed in IgA-expressing B-cells in the cecal tonsils of the birds **(Figure 3D)**. In contrast, we did not notice any changes in T-cell populations (CD3^+^, CD4^+^, and TCRγδ^+^) among the groups **(Figure S4, Supporting Information)**.

### Induction of strong cellular responses in immunised birds

A critical analysis suggests upregulation of major pro-inflammatory cytokines and transcription factors (IL-8, IL-1β, IL-17A, TNF-α, IFN-γ and NF-κB) in Splenic and BOF tissues of birds mucosally administered with multi-component vaccines (CadF, JlpA, and Hcp) compared to birds that received empty *L. lactis* NZ9000 cells **(Figure 4A, B)**.

**Figure 4:**
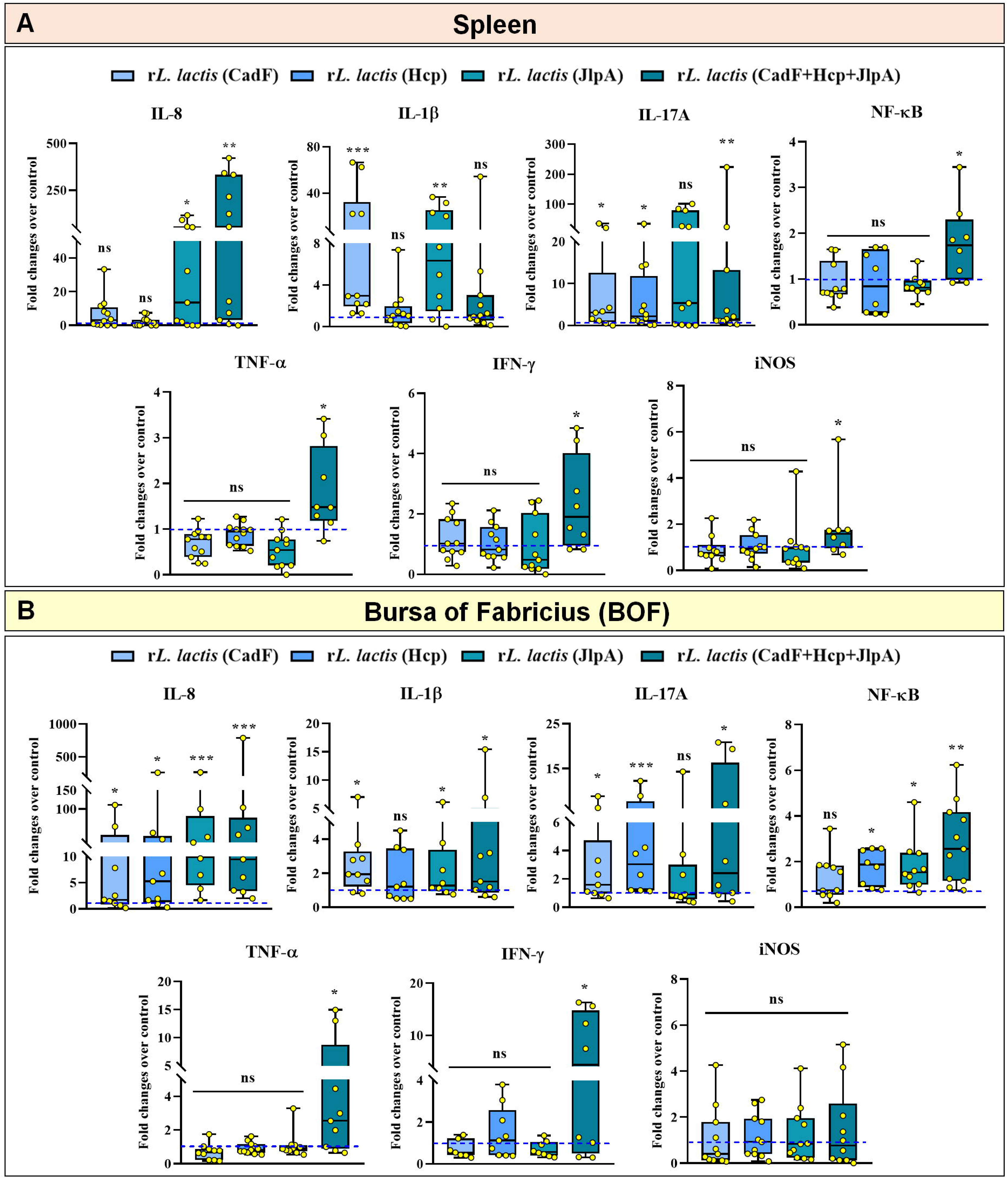
Cytokine gene expression profile in splenic and bursal tissues. **A:** Spleen tissues collected on day 28 post-first administration were subjected to mRNA extraction to analyse the gene expression profile using qRT-PCR. Data show significant upregulation of IL-8, IL-17A, TNF-α, NF-κB, IFN-γ and iNOS genes in birds that received the combination of all three proteins r*L. lactis* (CadF+Hcp+JlpA). Data represent the mean fold changes of mRNA expression over control (*L. lactis*) (dotted blue line) ± SE of 12 birds/group from two independent experiments (n =12). **B:** Bursa tissue was also collected on day 28, and mRNAs extracted from bursal tissue showed significant upregulation of IL-8, IL-1β, IL-17A, TNF-α, IFN-γ, and NF-κB genes in r*L. lactis* (Cad+JlpA+Hcp) group. Fold changes were calculated with respect to the control group (received WT *L. lactis* only). Data represent the mean fold changes of mRNA expression over control (*L. lactis*) (dotted blue line) ± SE of 12 birds/group from two independent experiments (n =12).

### Significant decrease in the cecal load of *C. jejuni* in birds

To assess the effect of mucosal administration of r*L. lactis* expressing CadF, Hcp and JlpA, the cecal load of each bird was measured on day 7 post-challenge with *C. jejuni*. The results indicate a significant reduction in the bacterial load in the cecum in immunised birds receiving the combination of all three proteins compared to those receiving individual proteins or the control group (immunised vs. control) **(Figure 5A)**.

**Figure 5:**
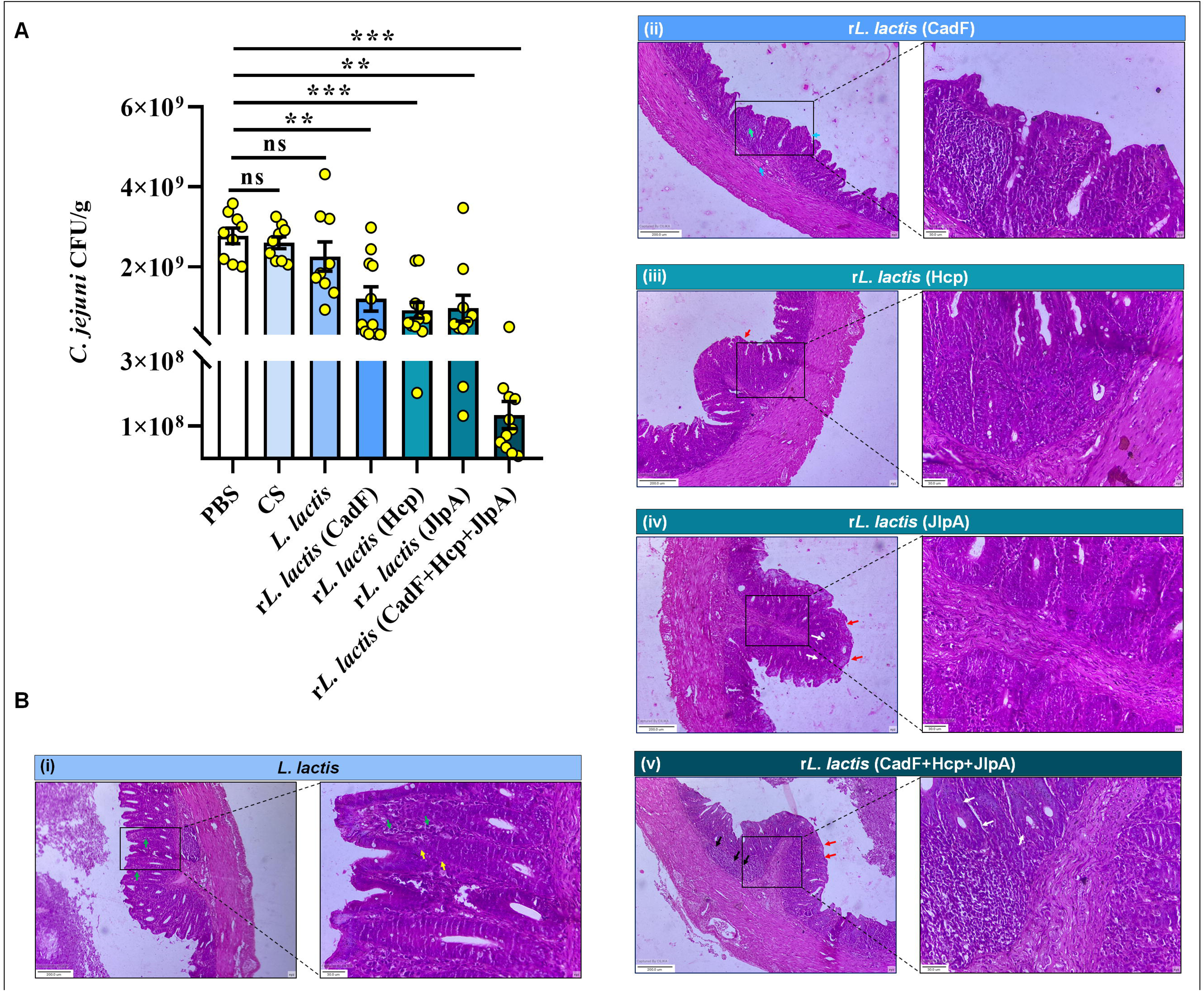
Reduction in cecal load of *C. jejuni* and histopathological analysis of cecal tissue. **A:** Birds administered with different combinations of r*L. lactis* were challenged with 1×10^8^ CFU *C. jejuni* TGH 9011. On day 7, post-challenge cecal contents were collected and processed to determine the cecal load of *C. jejuni*. Comparative analysis of bacterial load among different experimental groups shows a significant reduction in the birds that received r*L. lactis* expressing CadF, Hcp and JlpA (alone or in combination) compared to the control group (PBS only). Data represent the mean CFU/gm ± SE of two independent experiments (n = 12) performed under similar conditions. Asterisks indicate a statistically significant difference compared to the PBS group (***p ≤ 0.001). **B:** Histopathological analysis of formalin-fixed paraffin-embedded sections of the cecal tissue collected from birds at day 7 post-infection with *C. jejuni* TGH 9011. The different coloured arrows indicate outcomes in histopathology. Yellow arrow- necrosis, green arrow- infiltration, cyan arrow- reduced oedema, light green arrow- reduced infiltration, red arrow- intact epithelia, white arrow-well-structured crypt, black arrow-lymphoid accumulation.

### *C. jejuni* challenge in vaccinated birds develops little or no gut lesions compared to unvaccinated birds

Finally, tissue sections from each group were subjected to histopathological analysis to see the effect of vaccination on preserving gut health in cecal tissue against *C. jejuni* challenge infection. Birds that received the empty vector (*L*. *lactis* only) exhibited significant infiltration of pro-inflammatory cells, particularly heterophils and lymphocytes, within the cecal mucosa and submucosa. These birds also showed clear evidence of oedema and blood vessel congestion. The typical architecture of the cecum was severely disrupted, with signs of mucosal erosion, ulceration, and crypt distortion, which indicated compromised epithelial integrity **(Figure 5B-i)**.

In contrast, birds immunised with individual proteins or in combination demonstrated markedly improved gut health at post-challenge time points **(Figure 5B-ii, iii, iv)**. However, immunisation with CadF alone resulted in only mild mucosal erosion **(Figure 5B-ii)**. Notably, the combination vaccination with CadF, JlpA, and Hcp provided complete protection of the cecal tissue. Birds in this group displayed well-preserved cecal mucosal architecture, intact epithelial lining, organised crypts, and visible lymphoid aggregates. There was no evidence of mucosal erosion, ulceration, necrosis, or haemorrhage, suggesting that the combined vaccine offers the highest level of protection against *C. jejuni* challenge **(Figure 5B-v) (Figure S5, Supporting Information)**.

### Analysis of gut (cecal) microbial composition in immunised birds

#### Relative abundance of top ten phyla

Full-length 16S rDNA amplicon sequencing using the MinION™ nanopore long-read sequencer revealed significant variation in the cecal microbiota across different experimental groups. The taxonomic profiling at the phylum level within the top ten most abundant taxa reveals a predominance of three phyla: Bacillota (formerly *Firmicutes*), Pseudomonata and Bacteroidota. Bacillota remains the most dominant phylum, followed by Pseudomonata and Bacteroidota. Analysis of the top ten most abundant phyla revealed distinct patterns across experimental groups. Notably, Bacillota, a phylum commonly associated with gut health and metabolic homeostasis, showed consistently high abundance among all groups. Birds administered with r*L. lactis* expressing the Hcp antigen exhibited the highest average abundance of Bacillota (∼94.06%), compared to the group receiving empty vector (*L. lactis* only) (∼91.26%). Interestingly, Bacteroidota, another important phylum involved in carbohydrate fermentation and gut barrier integrity, was most abundant (∼2.33%) in birds that received a combination of all three recombinant *L. lactis* strains (expressing CadF, Hcp, and JlpA). This enrichment may indicate synergistic benefits of the present multivalent vaccine approach using probiotic bacteria on gut microbial balance.

In contrast, administration of the recombinant *L. lactis* cocktail (CadF+Hcp+JlpA) led to a ∼2- fold reduction in the relative abundance of Campylobacterota and Pseudomonata, two phyla known as pathogenic and opportunistic bacteria. However, no significant differences were observed across groups in the abundance of other major phyla, including Actinomycota, Cyanobacteriota, Spirochaetota, and Mycoplasmatota, indicating that the probiotic intervention did not broadly disrupt the existing gut microbial community structure **(Figure 6A, B) (Table S5, Supporting Information)**.

**Figure 6:**
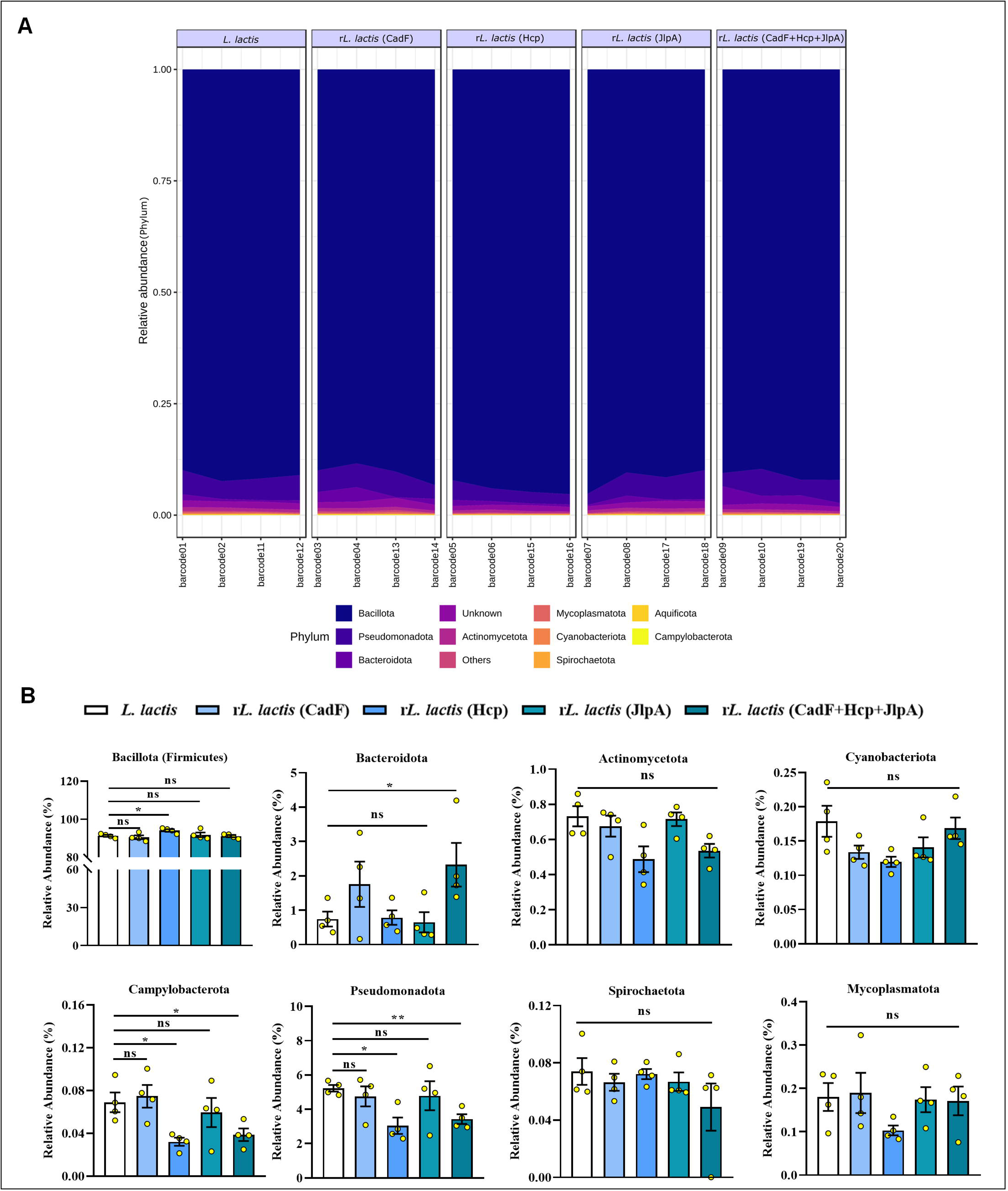
Relative abundance of cecal microbiota at the phylum level. **A:** The relative abundances of cecal microbiota at the phylum level in birds with different r*L. lactis* treatments based on full-length 16S sequencing are shown as stacked area plots, n=4 per group. **B:** The filtered count of the top 10 phyla in the cecal microbiota of different birds showed that Bacillota was the top phylum and showed steadily high abundance among all groups. Bacteroidota was abundant in birds that received a combination of all three recombinant *L. lactis* (expressing CadF, Hcp, and JlpA). r*L. lactis* combination led to a reduction in the relative abundance of Campylobacterota and Pseudomonata. No significant differences were observed across groups in the abundance of other major phyla, including Actinomycota, Cyanobacteriota, Spirochaetota, and Mycoplasmatota.

#### Relative abundance of top twenty genera

Further taxonomic analysis at the genus level revealed that *Lactobacillus* was the most dominant genus within the phylum Bacillota, with notable variation across experimental groups. In particular, birds immunised with r*L. lactis* expressing Hcp, as well as those receiving the combination of all three recombinant strains (CadF+Hcp+JlpA), exhibited the highest relative abundance of *Lactobacillus* (∼29%), in contrast to a lower abundance observed in the control group (∼17%). Other genera among the top 20 most abundant included *Blautia, Mediterraneibacter, Sellimonas, Streptococcus, Clostridioides,* and *Bacillus*. However, these genera did not exhibit substantial differences in abundance across experimental groups **(Figure 7A, B)**.

**Figure 7:**
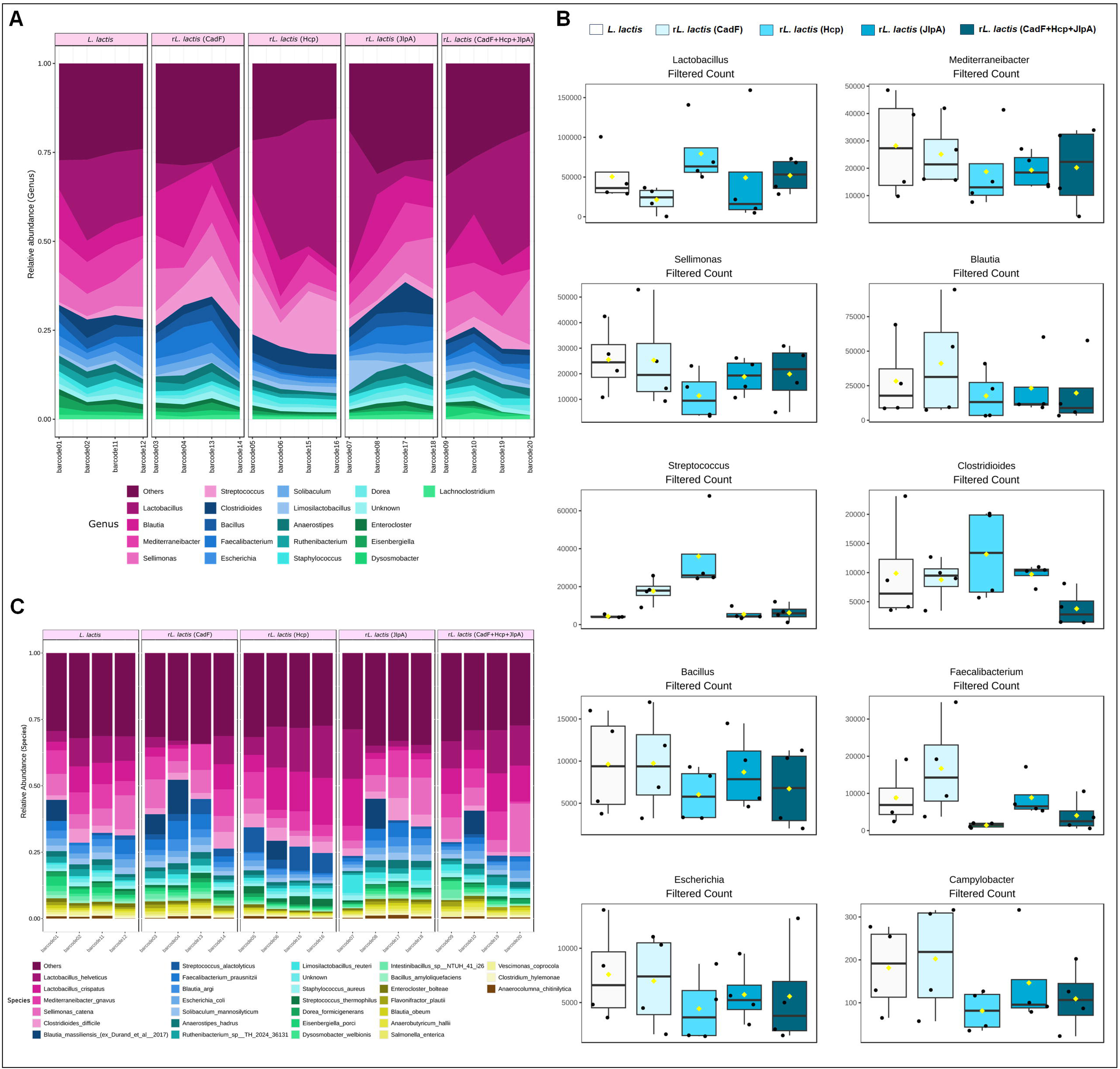
Relative abundance at the genus and species level of cecal microbiota. **A:** The relative abundances of the top 20 genera in the cecal microbiota of the birds based on full-length 16S rDNA sequencing are shown as stacked area plots. Birds immunised with *rL. lactis* expressing Hcp, and those receiving the combination of all three recombinant strains (CadF+Hcp+JlpA) exhibited the highest relative abundance of *Lactobacillus* compared to the control group. **B:** The filtered count of the top 10 genera in the cecal microbiota of different birds. **C:** The relative abundance of the top 30 species in the cecal microbiota of the birds is shown in stacked bar plots. *Lactobacillus helveticus* and *Lactobacillus crispatus* appeared to be the most abundant species in the Hcp and combined (CadF+Hcp+JlpA) group.

#### Relative abundance of top thirty species

At the species level, analysis of the top 30 most abundant taxa further clarified the specific microbial shifts associated with recombinant *L. lactis* immunisation. Within the dominant genus *Lactobacillus*, *Lactobacillus helveticus* and *Lactobacillus crispatus* emerged as the most abundant species, particularly in the Hcp- immunised and combined antigen (CadF+Hcp+JlpA) groups **(Figure 7C).**

### Alpha and beta diversity metrics of the cecal microbiota

#### Alpha diversity

Alpha diversity matrices (Chao1, Shannon, Simpson, and Fisher indexes): Chao1 estimates taxa richness by accounting for undetected features because of low abundance. Shannon and Simpson consider species richness and evenness, with varying weight given to evenness, and Fisher models the community abundance structure as a logarithmic series distribution. The alpha diversity was reduced in birds that received the r*L. lactis* (Hcp) treatment compared to the *L. lactis* and r*L. lactis* (CadF) groups by both Shannon and Simpson diversity measures. In Chao1 and Fisher indexes, no significant differences were observed among different treatment groups **(Figure 8A-D) (Table S6, Supporting Information)**.

**Figure 8:**
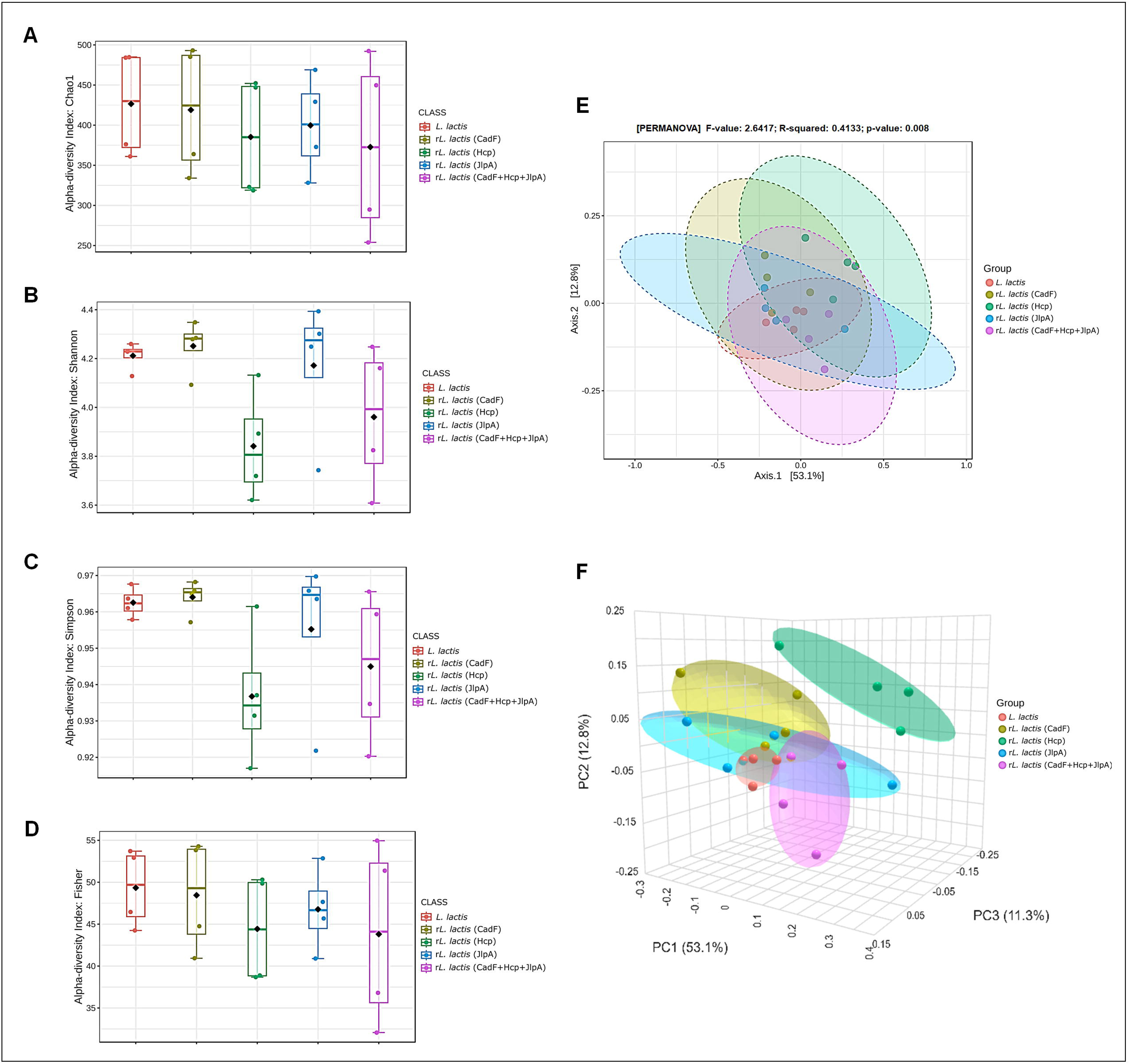
Alpha and Beta diversity matrices among groups. **A-D :** Alpha diversity matrices of the cecal microbiota of birds were evaluated using Chao1, Shannon, Simpson, and Fisher diversity indexes. The Chao1 index was used to assess the species richness. The microbial communities’ evenness and richness were both reflected in the Shannon index. The Simpson index highlighted dominance, and with less impact from evenness, Fisher’s alpha displayed the species richness. **E-F :** Beta diversity was assessed with the Bray-Curtis dissimilarity index and statistically tested by the PERMANOVA test (P < 0.05) and represented by the Principal Coordinates Analysis (PCoA) plot and 3D plot.

#### Beta diversity and community structure shifts

To assess overall differences in microbial community composition among the treatment groups, Principal Coordinates Analysis (PCoA) was performed based on Bray-Curtis dissimilarity. The resulting ordination plot revealed distinct clustering patterns among the experimental groups. Birds immunised with r*L. lactis* (Hcp) recorded significant shifts (p<0.05) in community structure compared to the control group (birds receiving the empty vector, *L. lactis* NZ9000) and r*L. lactis* (CadF+Hcp+JlpA) **(Figure 8E, F) (Table S7, Supporting Information)**.

### Differential abundance analysis of microbiota in birds

The LEfSe analysis showed distinct microbial shifts across the different *L. lactis* treatment groups. The r*L. lactis* (Hcp) group was enriched in taxa, such as *Streptococcus* and *Romboutsia*, which include species with recognised probiotic potential. However, this treatment also showed the most significant number of taxa depletions relative to *L. lactis* control, including genera associated with potential pathogenicity or pro-inflammatory activity, such as *Clostridium*, *Bacteroides*, *Vibrio*, *Pseudomonas*, and *Tropheryma*. In contrast, the r*L. lactis* (CadF+Hcp+JlpA) group uniquely enriched *Parabacteroides*, *Intestinimonas*, and *Flavonifractor*, all known producers of short-chain fatty acids **(Figure 9A)**.

**Figure 9:**
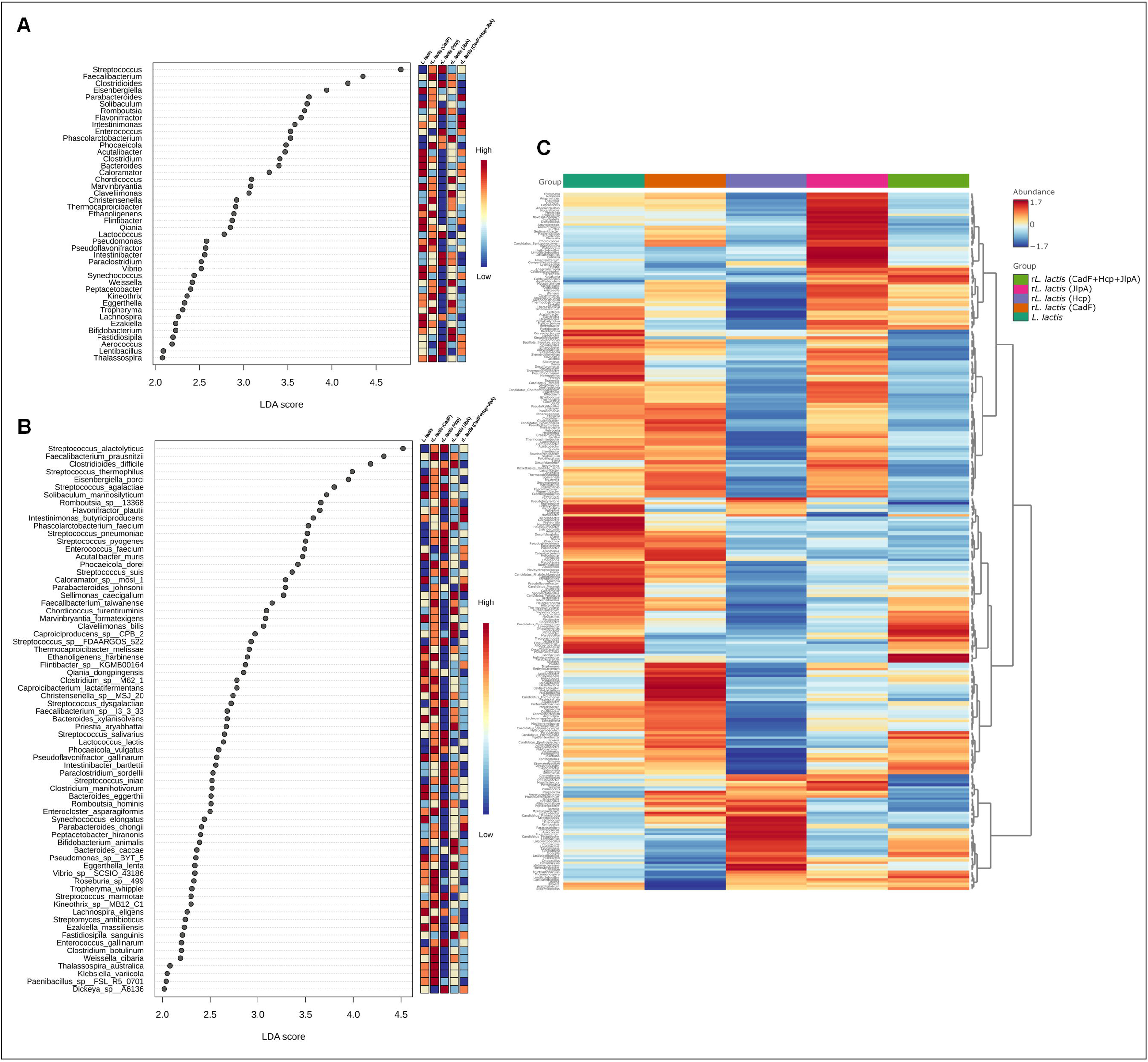
Differential abundance analysis and microbial cluster patterns among groups. **A-B:** Differential abundance analysis comparing bacterial taxa in the cecum of experimental birds with different r*L. lactis* treatments. The dot plot showed the results from LEfSe analysis, showing the significantly differential taxa at the genus and species level (LDA score >2, P<0.05) in the cecal microbiota of birds. **C:** The clustering heatmap showed the relative abundance of microbial genera between treatment groups.

At the species level, distinct microbial signatures were observed across the treatment groups, mirroring the genus-level patterns. The r*L. lactis* (Hcp) group exhibited the most extensive microbial depletion, including species such as *Thermocoproicibacter melissae*, *Streptomyces antibioticus*, and other fermentative taxa. Despite this, it uniquely enriched *Streptococcus* spp. and *Peptacetobacter hiranonis*, known secondary bile acid producers. In contrast, the r*L. lactis* (CadF+Hcp+JlpA) group selectively enriched short-chain fatty acid producers such as *Parabacteroides johnsonii*, *Intestinimonas butyriciproducens*, and *Flavonifractor plautii*. However, this group also showed depletion of potentially beneficial species, including *Peptacetobacter hiranonis* and *Streptococcus salivarius*, which are associated with bile acid metabolism and potential probiotic functions **(Figure 9B)**.

### Microbial composition and cluster patterns across treatment groups

The heatmap showed the relative abundance of microbial genera between treatment groups. The *L. lactis* group demonstrated a distinctive microbial signature with relatively intermediate abundance patterns for most taxa. The combined group exhibited the most distinctive microbial profile, characterised by a heavy enrichment or depletion of several genera relative to the *L. lactis* group. Significant divergence from the *L. lactis* group was also observed in the r*L. lactis* (Hcp) and r*L. lactis* (JlpA) groups, which formed separate clusters and displayed differential abundance patterns with selectively enriched red and depleted blue areas, indicative of selective enrichment or suppression of important bacterial taxa. Interestingly, the r*L. lactis* (CadF) group was positioned near the *L. lactis* in the microbial organisation but still displayed taxa-specific changes. Group-specific changes in abundance were observed in species such as *Lactobacillus, Clostridium, Ruminococcus, and Bacteroides*, where specific genera are steadily enriched or depleted through recombinant treatments, highlighting the impact of antigen- specific immune modulation on gut microbial dynamics **(Figure 9C)**.

## DISCUSSION

Together with the emergence of new drug-resistant strains of *C. jejuni* and their intrinsic ability to colonise the chicken gut pose substantial risks to food safety and public health [54–56]. On the other hand, vaccination against *C. jejuni* remains challenging due to several key factors, including its high genetic diversity, frequent phase variation, extensive serotype diversity, and immune evasion [57–59]. Given that *C. jejuni* colonisation is a multifactorial process, one approach to address these challenges is to design a multi-component vaccine using immunogenic subunits with high sequence conservation across diverse *C. jejuni* strains. To this end, several surface-exposed and secretory proteins of *C. jejuni* have been identified as potential vaccine targets due to their sequence conservation and roles in bacterial adherence, invasion, and virulence expression. Among these proteins, Hcp, JlpA, CadF, and FlpA, along with other major outer membrane proteins (MOMPs) like PorA, CmeC, and PEB1, have emerged as promising vaccine candidates for controlling *C. jejuni* infections [59–64]. Specifically, the *C. jejuni* CadF and JlpA proteins are two highly conserved key SECPs mediating host cell adherence by binding to fibronectin and the tissue chaperone Hsp90α, respectively. These interactions represent crucial early steps in bacterial colonisation to trigger pro-inflammatory host responses [16,65–67].

On the other hand, as a critical effector protein of the functional Type VI secretion system (T6SS), we recently reported that *C. jejuni* Hcp can contribute to bacterial competition and environmental adaptation. In addition, we also showed that Hcp enhances competitive fitness in a dynamic gut environment, and mucosal delivery of recombinant Hcp can drive robust immune responses within the intestinal mucosa in avian and murine models [17,49]. Thus, with its dual role in pathogenesis and immunogenicity, the high sequence conservation of *hcp* among *C. jejuni* strains makes it another compelling vaccine candidate. Moreover, our *in-silico* studies could identify multiple conserved putative B-cell epitopes within the sequence of CadF, Hcp and JlpA, strengthening their potential as promising multi-component vaccine targets to limit the cecal load of *C. jejuni* in chickens [17,18].

To this end, the present study aims to evaluate the prospective benefits of a multivalent vaccine strategy, incorporating three immunogenic proteins (CadF, Hcp and JlpA) with distinct functionality to determine whether a multivalent formulation can offer superior protection compared to a monovalent approach.

For this purpose, we used bioengineered recombinant *L. lactis* surface-expressing CadF, Hcp, and JlpA proteins of *C. jejuni*. To enhance mucosal adherence and intragastric stability and prolong gut transit time, we further coated the recombinant bacteria with chitosan. As a natural biopolymer, chitosan is a linear cationic polysaccharide consisting of N-acetyl-d-glucosamine and d-glucosamine coupled by β (1→4) glycosidic linkages. This conformation enables chitosan to bind sialic acid and other negatively charged biomolecules in mucus and epithelial cell surfaces through electrostatic interactions. Additionally, incorporating sodium tripolyphosphate (STPP), a non-toxic polyanionic salt, facilitates cross-linking of chitosan, thereby enhancing the gelation and reticulation capacity of the resulting biopolymer [68–71]. Previously, we showed that CS-TPP coating of the LAB vector does not adversely affect bacterial growth; rather, it acts as a protective barrier against a range of physicochemical gut-mimicking conditions [37]. We confirmed that the CS coating of *L. lactis* cells does not interfere with the accessibility of CadF, Hcp, and JlpA proteins expressed on the bacterial surface. Furthermore, *in vitro* cell adhesion studies supported our hypothesis that chitosan- coating enhances the bacterial ability for cell association due to its strong mucoadhesive properties [72–74]. This observation aligns with our previous findings, which demonstrated that chitosan coating can extend the gut transit time of LAB vectors in both murine and avian models [37]. This has remained a strong foundation for us to explore the *in vivo* application of non-commensal LAB strains, such as *L. lactis*, as live vector vaccine delivery platforms against *C. jejuni.* We showed that oral administration of chitosan (CS)-coated recombinant *L. lactis* expressing the three antigenic proteins (CadF, Hcp, and JlpA) resulted in a significant increase in local antibody levels, specifically secretory IgA (sIgA) against *C. jejuni* whole-cell antigens in both intestinal lavages and freshly collected faecal pellets. Compared to immunisation with individual antigens, enhanced mucosal antibody responses by the combined delivery of all three subunits suggest the more pronounced effect of the “combinatorial approach” at the mucosal surface. In contrast, oral administration of r*L. lactis* did not influence systemic antibody (IgY) levels, likely due to the limited trafficking of mucosally activated IgY- producing B cells into the systemic circulation [18,20]. Nevertheless, both control groups (PBS and CS) of birds failed to induce secretory IgA; therefore, apart from cecal load determination, we did not include them in further experiments. Instead, comparisons were made with the CS-coated WT *L. lactis*. Importantly, we also demonstrated that a cocktail of antibodies harvested from intestinal lavages neutralised *C. jejuni* adherence and invasion of primary CEICs. These findings further confirm that local antibodies raised by mucosal delivery of r*L. lactis* expressing CadF, Hcp, and JlpA can specifically recognise and bind to their corresponding surface antigens on *C. jejuni*, thereby interfering with bacterial colonisation and invasion mechanisms.

Since vaccine efficacy depends on the induction of both humoral and cellular immune responses, we evaluated the cellular immune response in peripheral lymphoid organs to assess the potential of the current vaccine formulation in promoting long-term immunity, memory formation, and the orchestration of antibody production. To this end, our immunophenotyping data indicate some synergistic effect in IgA^+^ B cell populations in the Bursa of Fabricius (BOF) in birds that received the combination of all antigens compared to those immunised with a single antigen. However, no noticeable changes were observed in the CD3[, CD4[, and TCRγδ[T cell populations in the cecal tonsils of immunised birds **(Figure S4, Supporting Information)**. This may be due to the nature of the current mucosal vaccine modality using recombinant LAB vectors, which appears to be less effective at inducing T cell differentiation and instead preferentially activates B cells to provide immediate mucosal protection. While the humoral and cellular arms of the immune system are inherently interconnected, our findings further highlight a synergistic enhancement of cellular immunity in birds vaccinated with the combined antigen formulation (CadF, Hcp, and JlpA). Specifically, these birds exhibited a marked splenocyte proliferative response and increased nitric oxide (NO) production upon exposure to *C. jejuni* whole-cell lysate, indicating effective activation of immune effector functions. Notably, the increased proliferation and NO production correlated with concurrent upregulation of key pro-inflammatory and immune-regulatory genes, including IL-8, IL-1β, IL- 17A, TNF-α, and NF-κB, in birds that received r*L. lactis* expressing CadF, JlpA, and Hcp either individually or in combination. These findings suggest that recombinant *L. lactis*-based mucosal immunisation, particularly with multiple antigens, can elicit both innate and Th1-type systemic immunity. The involvement of NF-κB, a central transcription factor in inflammation and host defence, further supports the immunostimulatory potential of this vaccine strategy [75,76].

Finally, to determine whether a Th1-biased response and potent neutralising antibodies can restrict bacterial colonisation, we performed an *in vivo* challenge with highly pathogenic *C. jejuni* (TGH 9011) in immunised birds. The results from our *in vivo* challenge experiment strongly support the efficacy of the multivalent vaccine composition in limiting bacterial colonisation in the intestine **(Figure 5A)**. Moreover, histopathological analysis of the cecal tissue of vaccinated birds challenged with *C. jejuni* exhibited well-preserved intestinal architecture, with intact villi, minimal infiltration of inflammatory cells, and reduced epithelial damage, compared to the unvaccinated controls, which showed marked villus blunting, mucosal erosion, and lymphocytic infiltration **(Figure 5B)**. Together, these findings suggest that the multivalent vaccine reduced pathogen burden, mitigated gut inflammation, and maintained mucosal integrity.

Notably, irrespective of the experimental groups, a reduction in *C. jejuni* cecal load was observed, presumably due to a non-specific probiotic effect and the addition of *L. lactis* to each treatment regimen. In particular, several *L. lactis* strains, including the one used in this study, have demonstrated mucus-binding properties [77,78]. These properties are associated with beneficial gut-targeted activities that may enhance microbial fitness and persistence within the host gastrointestinal tract [79].

Vaccination can significantly impact poultry health both positively and, in some cases, negatively, depending on the route of administration, vaccine composition, dosage, and vaccination regimen. In commercial poultry production systems, the vaccination approach must aim to enhance immunoprotection and ensure improved productivity, particularly in terms of growth rate, feed conversion efficiency, and egg production. Given these considerations, we selected probiotic bacteria as a delivery platform for the present vaccine formulation, anticipating that this approach would offer dual benefits: enhanced immune protection and improved gut health. Given that vaccine-induced modulation of the gut microbiota can influence immune signalling pathways and vice versa, we further examined the microbial composition and diversity in the cecal contents of the immunised birds.

The analysis of full-length 16S rDNA sequencing of the cecal microbial community highlighted the predominance of Bacillota (formerly *Firmicutes*) and Bacteroidota, suggesting that immunisation may have modulated the gut microbiota in a favourable direction, contributing to improved gut health, as corroborated by histopathological findings. In particular, the elevated abundance of Bacillota and Bacteroidota in the immunised group points to a possible selective expansion of beneficial microbial populations. Additionally, a significant reduction in the relative abundance of Pseudomonodota and Campylobacterota in either the r*L. lactis* (Hcp) or the combined treatment group indicates depletion of the harmful microbiota. These findings support the notion that the administration of recombinant *L. lactis* strains may exert a protective effect by promoting the selected growth of beneficial microbes while suppressing potentially harmful populations **(Figure 6A, B)** [80–82].

Further taxonomic analysis at the genus level revealed that *Lactobacillus* was the most dominant genus within the Bacillota phylum, with its abundance showing notable variation across experimental groups. The enrichment of *Lactobacillus*, a well-characterised probiotic genus known for enhancing intestinal barrier integrity, competitively excluding pathogens, and modulating host immune responses, reinforces the health-promoting potential of recombinant *L. lactis*, especially when engineered to express immunogenic antigens such as Hcp **(Figure 7A, B)**.

When searching for other top genera, *Blautia, Mediterraneibacter, Sellimonas, Streptococcus, Clostridioides,* and *Bacillus* are of note; however, no noticeable changes could be observed **(Figure 7A, B)**. This led us to interpret that the *L. lactis*-mediated influence on gut microbial community structure was selective rather than broad-spectrum, likely driven by specific probiotic-host or probiotic-microbe interaction. To further assess the beta diversity and shifts in community structure, the resulting ordination plot revealed distinct clustering patterns among the experimental groups. Specifically, birds that received the combination of r*L. lactis* expressing the CadF, Hcp, and JlpA proteins exhibited broad dispersion along Axis 1 and 2. In contrast, birds immunised with r*L. lactis* (Hcp) recorded significant shifts in community structure compared to the control and combined groups, formed relatively tighter and more coherent clusters, suggesting limited diversity in microbial communities **(Figure 8E, F)**. Moreover, differential abundance analysis showed that the combined antigen formulation (CadF, Hcp, and JlpA) selectively enriches short-chain fatty acid producers. These distinct attributes in microbial community structure, along with data from the correlation network analysis of cecal microbiota **(Figure S6, Supporting Information)**, indicate that the effect of the *L. lactis*- mediated combined expression and delivery of immunogenic subunits of *C*. *jejuni* in the gut may synergistically enhance cross-taxa correlations, leading to profound, complex, and potentially beneficial changes in the gut microbial community of chickens.

## CONCLUSIONS

Our preliminary data, combined with the intrinsic adjuvanticity of the proposed vaccine strategy, underscore its potential in preserving gut homeostasis and safeguarding the intestinal mucosa against *C. jejuni* pathogenesis without inducing immune tolerance. Notably, bioengineering probiotic bacteria emerge as a promising live vector-based mucosal vaccine platform against common enteric pathogens, including *C. jejuni*, due to their capacity to stimulate both mucosal and, to some extent, systemic immune responses through the controlled expression of target proteins. Given that the efficacy of mucosal vaccines largely relies on regulating bacterial adhesion and invasion of the intestinal epithelium, optimal protein expression in the complex gut environment is critical to maximising the adaptability and effectiveness of LAB-based vaccine strategies.

## ASSOCIATED CONTENT

### Supporting Information

**Table S1:** The list of bacterial strains and plasmids, **Table S2:** Composition of poultry feed used in the present study **Table S3:** Detailed list of the antibodies used for flow cytometry, **Table S4:** Cytokine gene primers used for qRT-PCR**, Table S5:** Multiple linear regression with covariate adjustment of cecal microbiota at the phylum level, **Table S6:** Post-hoc pairwise comparison (multi-group) of alpha diversity indexes, **Table S7:** Pairwise PERMANOVA analysis of beta diversity, **Figure S1:** Bioengineering *L. lactis* surface expressing CadF protein of *C. jejuni*, **Figure S2:** Standard plot for sodium nitrite (NaNO_2_) using Griess reagent, **Figure S3:** *In vitro* adhesion assay of uncoated and CS-coated r*L. Lactis* (CadF, Hcp, JlpA) in CEICs**, Figure S4:** Flow cytometric analysis of T-cell subsets in cecal tonsils of immunised birds, **Figure S5:** Additional images of tissue sections showing histopathological changes of cecal tissue collected from birds at day 7 post-infection with *C. jejuni* (TGH 9011), **Figure S6:** Correlation network analysis of cecal microbiota.

## Abbreviations

LMICs: low- and middle-income countries
SECPs: surface-expressed colonisation proteins
LAB: lactic acid producing bacteria
CEICs: chicken embryo intestinal cells
RT: room temperature
STPP: sodium tripolyphosphate
RIR: rhode island red
BOF: bursa of fabricius
WCL: whole cell lysate
NO: nitric oxide
EDU: 5-Ethynyl-2′-deoxyuridine
TMB: 3,3’,5,5’-tetramethylbenzidine
Con A: concanavalin A
PFA: paraformaldehyde
CAT: cefoperazone, amphotericin B, and teicoplanin
H&E: hematoxylin and eosin
OTUs: operational taxonomic units
SparCC: sparse correlations for compositional data
MOMPs: major outer membrane proteins
T6SS: type VI secretion system.

## Authors Contributions

PB designed and performed the experiments, analysed the data and wrote the manuscript. SA helped with *in vivo* experiments and writing the draft. SM performed the histopathological analysis. SO helped in metagenomics data analysis. OG contributed to the data analysis and draft preparation. AIM conceived the study, analysed the data, and wrote the manuscript. All authors reviewed the manuscript.

## Acknowledgements

PB thanks CSIR, MoE, Govt. of India for the fellowship support. SA thanks UGC, MoE, Govt. of India for the fellowship support. AIM and OG thank BactiVac Network, UK, for funding this work. We acknowledge the DBS Central Imaging Facility, Flow cytometry facility and sequencing facility of IISER Kolkata. We thank IISER Kolkata for the immunofluorescence microscopy, flow cytometer and qRT-PCR facilities. We especially thank Mr. Tamal Ghosh from IISER Kolkata for handling the flow cytometry instrument and data analysis.

## DECLARATIONS

### Ethics approval and consent to participate

The chicken experimentation protocol was approved by the Institutional Animal Ethics Committee (IAEC), IISER Kolkata. All procedures were conducted following the guidelines of the Committee for Control and Supervision of Experiments on Animals (CCSEA), the Ministry of Fisheries, Animal Husbandry and Dairying, Government of India. The permit number for the *in vivo* chicken experiment is IISERK/IAEC/AP/2023/108, and for using chick embryos is IISERK/IAEC/AP/2023/107.

All procedures followed the institutional guidelines with prior approval from the Institute Biosafety Committee (IBSC), IISER Kolkata. The IBSC approval number for using *Campylobacter jejuni* is IISERK/IBSC/2019/02, and the *Lactococcus lactis* is IISERK/IBSC/2019/022.

### Consent for publication

Not applicable

### Availability of data and materials

The authors confirm that the data supporting the findings of this study are available within the article and its supplementary materials.

---

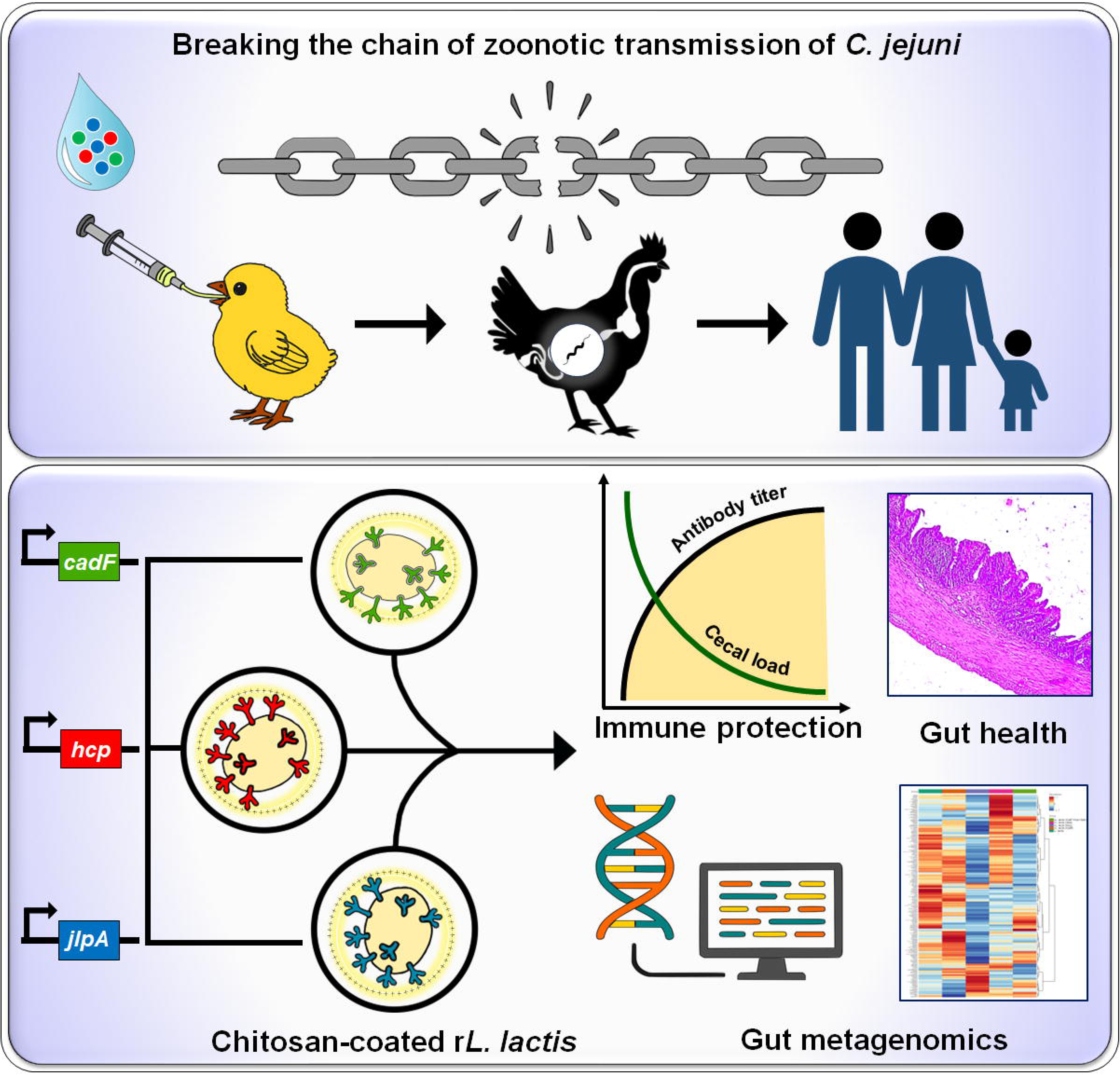

