## Supporting Information for "Recombinant *Lactococcus lactis*-based multivalent vaccine targeting *Campylobacter jejuni* colonisation and modulating cecal microbiota in poultry"

***Corresponding Authors**

**Dr. Ozan Gundogdu, PhD**

Associate Professor, Department of Infection Biology, Faculty of Infectious & Tropical Diseases, London School of Hygiene and Tropical Medicine, London, WC1E 7HT, United Kingdom, ORCID ID: 0000-0002-3550-0545.

| **Bacterial strains and**  **Plasmids** | **Characteristics** | **Purpose** | **Source** |
| --- | --- | --- | --- |
| **Strains** | | | |
| *E. coli* Top 10 | F- mcrAΔ (mrr-hsdRMS-mcrBC)  Φ80lacZΔM15 Δ lacX74 recA1  araD139 Δ (araleu)7697 galUgalKrpsL  (StrR) endA1 nupG | Recombinant plasmid  storage (*L. lactis-based*  plasmid constructs) | Thermo Fisher  Scientific, USA |
| *L. lactis* NZ9000 | MG1363 (nisRK genes into  chromosome), Wild type, plasmid-free | Wild-type control | Dr. L. G. Bermúdez  Humarán, INRA, France |
| *L. lactis* NZ9000- JlpA | Cm^r^, NZ9000 harboring pNZ8048-  CWAM6-JlpA | Recombinant protein  expression | ^1^ |
| *L. lactis* NZ9000- Hcp | Cm^r^, NZ9000 harboring pNZ8048-  CWAM6-Hcp | Recombinant protein  expression | ^2^ |
| *L. lactis* NZ9000- CadF | Cm^r^, NZ9000 harboring pNZ8048-  CWAM6-CadF | Recombinant protein  expression | This work |
| *C. jejuni* TGH9011 | T6SS-positive *C. jejuni* | *In vivo* challenge study | BEI Resources |
| **Plasmids** | | | |
| pNZ8048-CWA_M6_ | Cm^r^, SPUsp45, M6 cell wall anchor  expressed under the PnisA promoter | *L. lactis-*based recombinant  protein expression vector | Dr. L. G. Bermúdez  Humarán, INRA, France |
| pNZ8048-CWA_M6_-JlpA | Cm^r^, SPUsp45, M6 cell wall anchor  expressed under the PnisA promoter | *L. lactis*-based JlpA protein  expression vector | ^1^ |
| pNZ8048-CWA_M6_-Hcp | Cm^r^, SPUsp45, M6 cell wall anchor  expressed under the PnisA promoter | *L. lactis*-based Hcp protein  expression vector | ^2^ |
| pNZ8048-CWA_M6_-CadF | Cm^r^, SPUsp45, M6 cell wall anchor  expressed under the PnisA promoter | *L. lactis*-based CadF protein  expression vector | This work |

**Table S1:** The list of bacterial strains and plasmids.

| **Ingredients** | **Pre-starter feed (0 to 10 days)** | **Starter feed (11 to 35 days)** |
| --- | --- | --- |
| Maise | 550 | 590 |
| Soya Deoiled cake* | 390 | 340 |
| Oil | 16.5 | 27 |
| Dicalcium Phosphate | 12 | 12 |
| Line Stone Powder | 16 | 16 |
| Trace Mineral (Manganese, Zinc, Iron, Iodine, Copper, Cobalt) | 2 | 2 |
| Salt | 2.5 | 2.5 |
| Sodium bicarbonate | 1.5 | 1.5 |
| Choline Chloride | 0.5 | 0.5 |
| Lysine | 2.5 | 2 |
| D.L. Methionine | 2.8 | 2.7 |
| Toxin Binder | 1 | 1 |
| Emulsifier | 0.25 | 0.25 |
| Threonine | 0.15 | 0.15 |
| Phytase 5000 | 0.1 | 0.1 |
| Total | 999.9 | 999.8 |

**Table S2:** Composition of poultry feed used in the present study ^3^.

| **Antibody name** | **Fluorophore** | **Source** | **Volume utilised** |
| --- | --- | --- | --- |
| Anti-chicken Bu1 | FITC | Southern Biotech, Canada | 0.5 µL |
| Anti-chicken IgA | PE | Southern Biotech, Canada | 0.5 µL |
| Anti-chicken CD3ζ | APC | Southern Biotech, Canada | 0.5 µL |
| Anti-chicken CD4 | FITC | Southern Biotech, Canada | 0.5 µL |
| Anti-chicken TCR γδ | PE | Southern Biotech, Canada | 0.5 µL |
| 7-AAD | Far-red | Southern Biotech, Canada | 1 µL |

**Table S3:** Detailed list of the antibodies used for flow cytometry.

| **Primers** | **Sequence (5’→3’)** | **References** |
| --- | --- | --- |
| β-actin FP | GAGAAATTGTGCGTGACATCA | ^4^ |
| β-actin RP | CCTGAACCTCTCATTGCCA |  |
| CXCL8 FP | GGCTTGCTAGGGGAAATGA | ^5^ |
| CXCL8 RP | AGCTGACTCTGACTAGGAAACTGT |  |
| IL-17 FP | ATTCCAGGTGCGTGAACTCGGC | ^6^ |
| IL-17 RP | GTGCAGCCCACGGTGATCATTTTC |  |
| IFNγ FP | ACACTGACAAGTCAAAGCCGCACA | ^7^ |
| IFNγ RP | AGTCGTTCATCGGGAGCTTGGC |  |
| NFκB FP | GAAGGAATCGTACCGGGAACA | ^8^ |
| NFκB RP | CTCAGAGGGCCTTGTGACAGTAA |  |
| TNFα FP | TTGCAGGCTGTTTCTGCCTCTGC | This work |
| TNFα RP | TGAAGGTGGTGCAGATGGGGC |  |
| IL-1β FP | AGTGAGGCTCAACATTGCGCTGTA | ^9^ |
| IL-1β RP | TAGAAGATGAAGCGGGTCAGCTCG |  |
| iNOS FP | GAACAGCCAGCTCATCCGATA | This work |
| iNOS RP | CCCAAGCTCAATGCACAACTT |  |
| CadF FP | CGGCGCGCATGCGCTGATAACAATGTAAAATTTG | This work |
| CadF RP | CGCGCGGCTAGCGATCTTAAAATAAATTTAGCATCC |  |

**Table S4:** Cytokine gene primers used for qRT-PCR and CadF cloning primers.

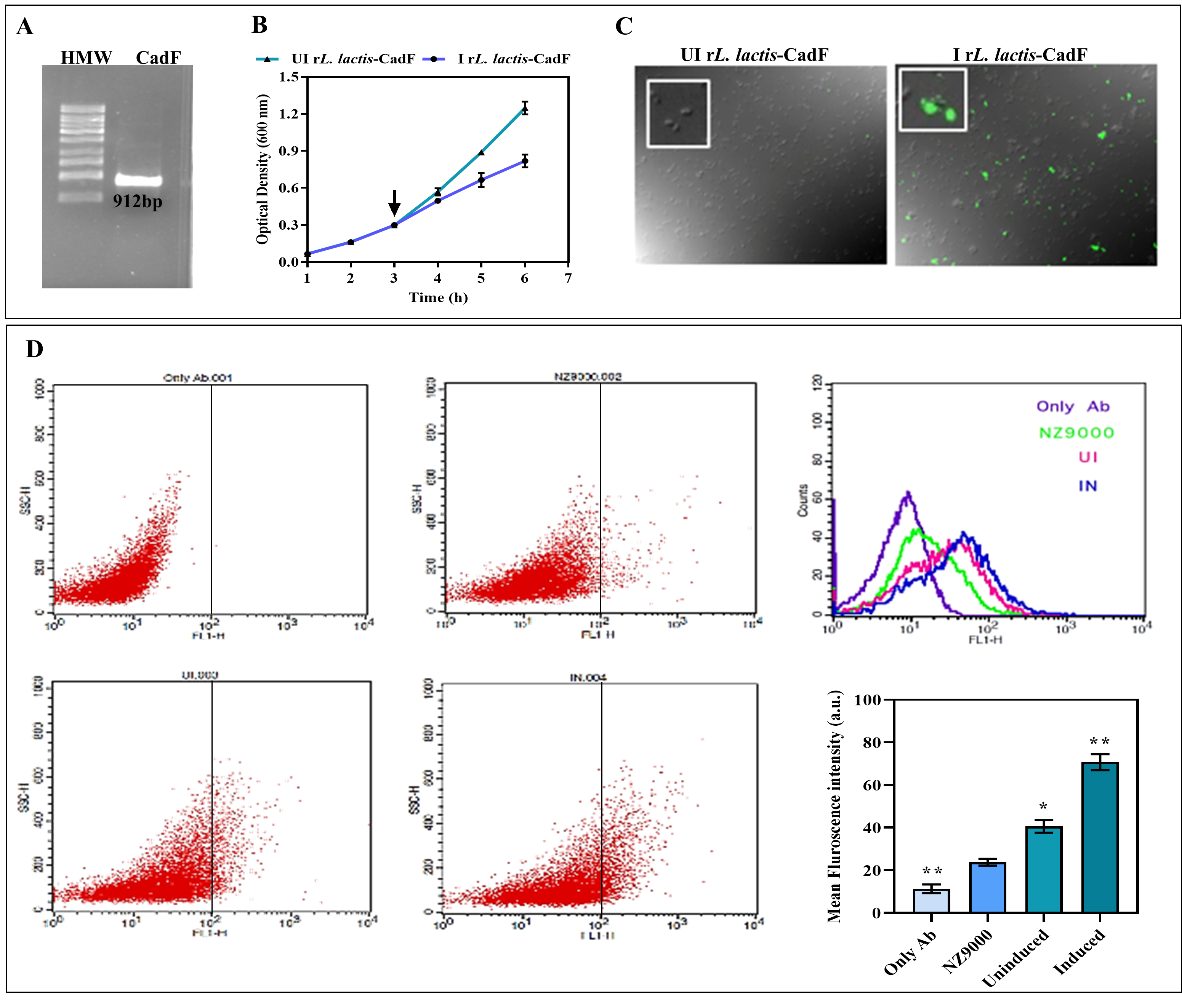

**Figure S1:** **Bioengineering *L*. *lactis* surface expressing CadF protein of *C. jejuni****.* The gene encoding the truncated CadF sequence was successfully cloned into the nisin-inducible *Lactococcal* expression vector pNZ8048-CWA_M6_, positioned between the N-terminal signal peptide (SP) of USP_45_ and the C-terminal CWA_M6_ domain. Restriction digestion of recombinant plasmid (pNZ8048-USP_45_**-CadF**-CWA_M6_) with *Sph*I/*Nhe*I endonucleases and gene sequencing confirmed the successful cloning of the desired gene. (**A)** *L. lactis* NZ9000 cells were electro-transformed with the pNZ-CadF plasmid, and the positive transformants were screened using chloramphenicol as a selection marker and gene amplification (~912 bp) using a specific primer set. (**B)** *In vitro* growth kinetics of uninduced and nisin-induced r*L. lactis* cells. Cells were induced with Nisin (Sigma, USA) (15 ng/mL) when OD_600_ reached ~0.2-0.3. **(C)** Further indirect immunofluorescent assay and (**D**) flow cytometric analysis of nisin-induced r*L. lactis* cells confirm optimal surface expression of CadF protein.

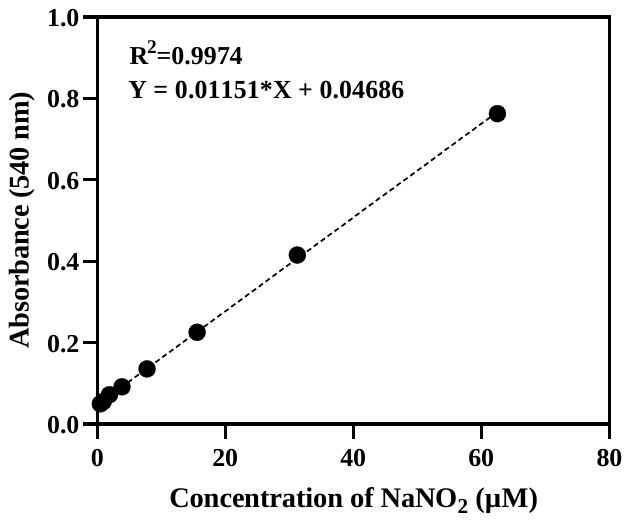

**Figure S2:** Standard plot for sodium nitrite (NaNO_2_) using Griess reagent. This plot was used to quantify NO production (refer to Figure 2D).

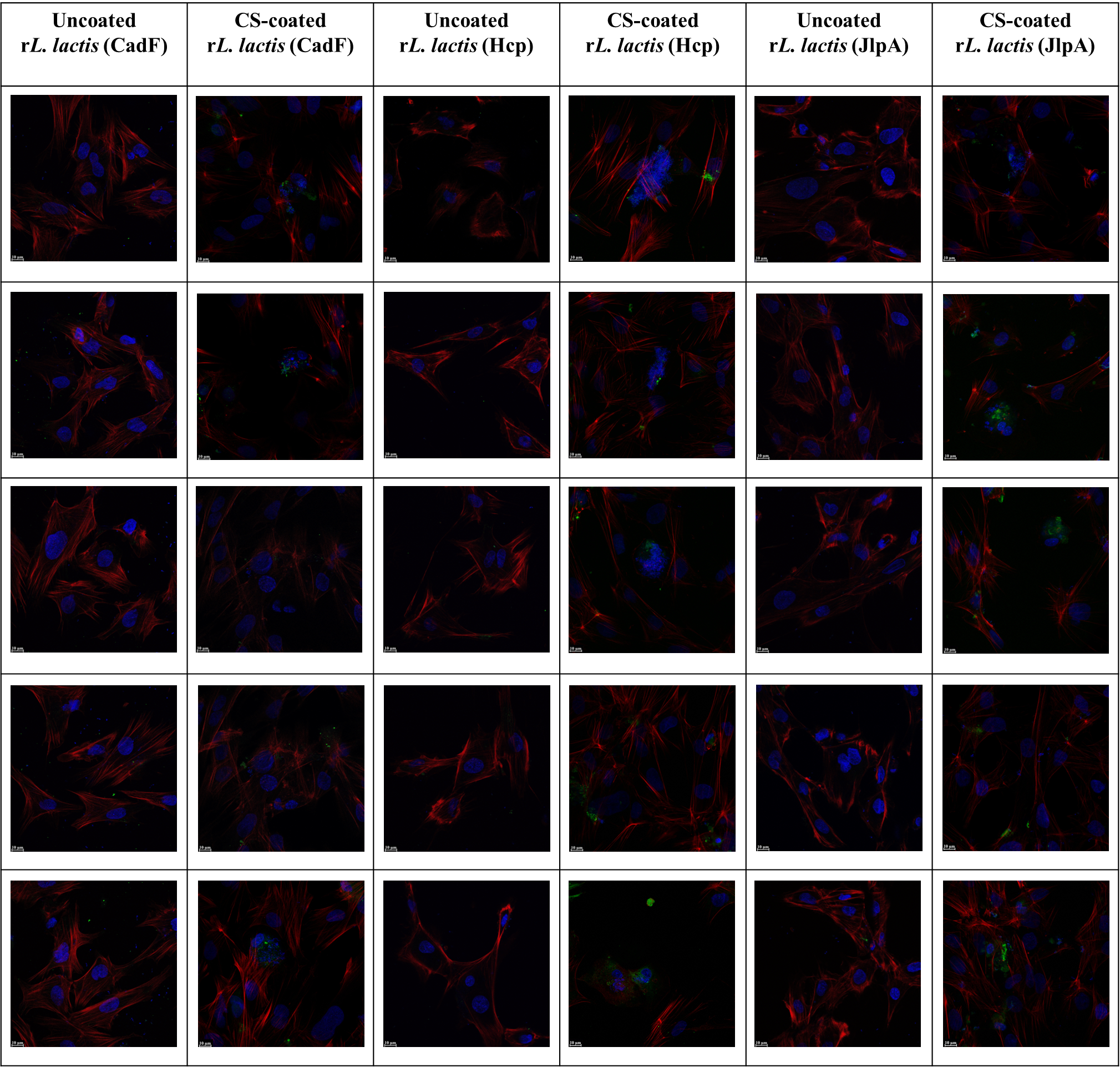

**Figure S3:** ***In vitro* adhesion assay of uncoated and CS-coated r*L. Lactis* (CadF, Hcp, JlpA) in CEICs.** The cells were incubated with induced uncoated and CS-coated r*L. lactis* 4 h at 37 °C with 5% CO_2_, at a ratio of cell vs bacteria 1:1000. CLSM images of cells incubated with CS-coated r*L. lactis* cells showed enhanced cell adhesion compared to uncoated bacteria. Scale bar: 10 μm.

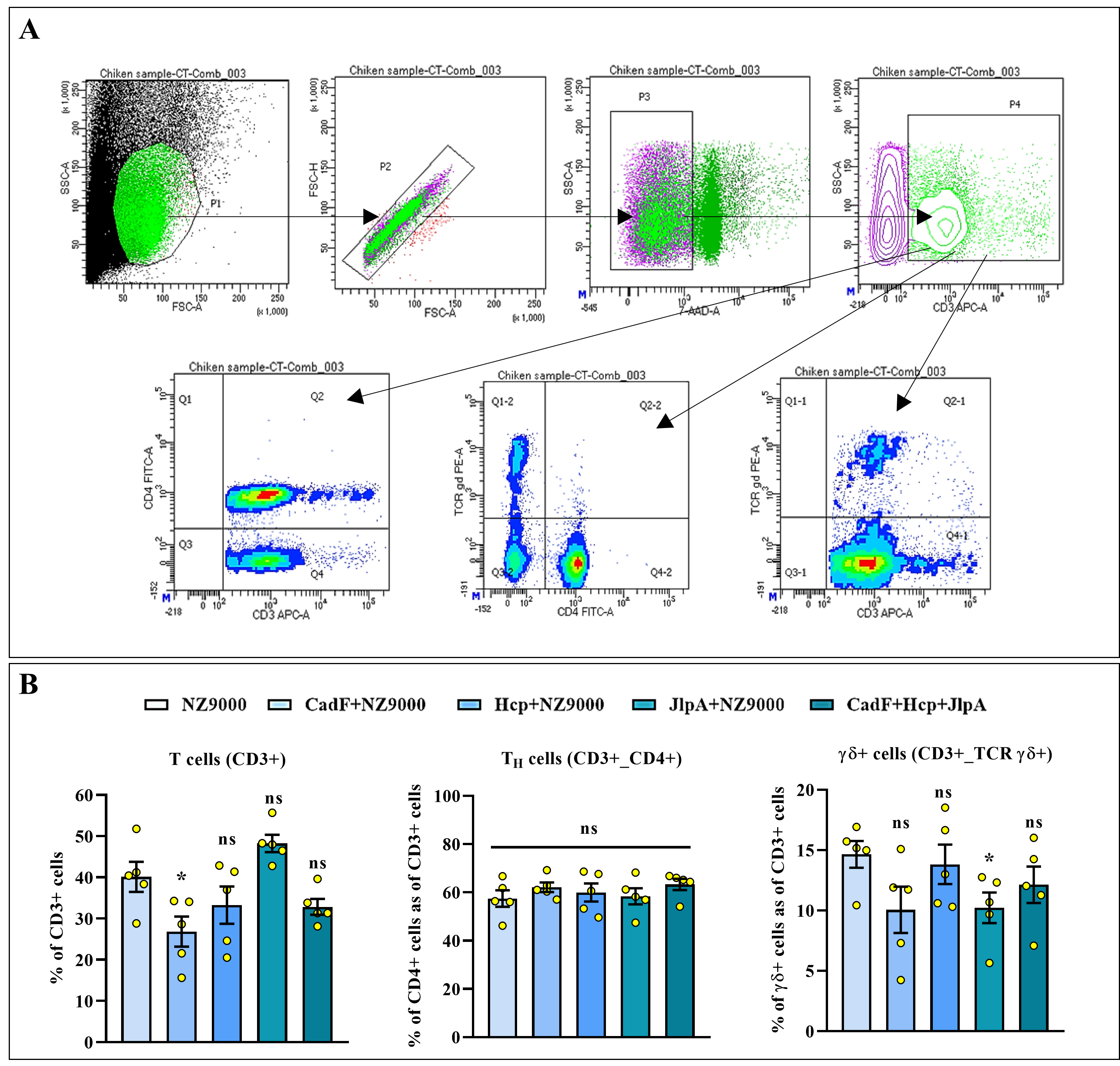

**Figure S4: Flow cytometric analysis of T-cell subsets in cecal tonsils of immunised birds.** Cecal tonsils were collected from birds at day 28 post-first immunisation and processed to isolate mononuclear cells. **(A)** Panel showing gating strategy for T cells present in the cecal tonsil (CT) of experimental birds. **(B)** Flow cytometric analysis of T cells suggests no changes in the CD3^+^, CD4^+^ and TCRγδ^+^ populations in birds administered with r*L. lactis* (CadF, JlpA and Hcp) compared to other experimental groups.

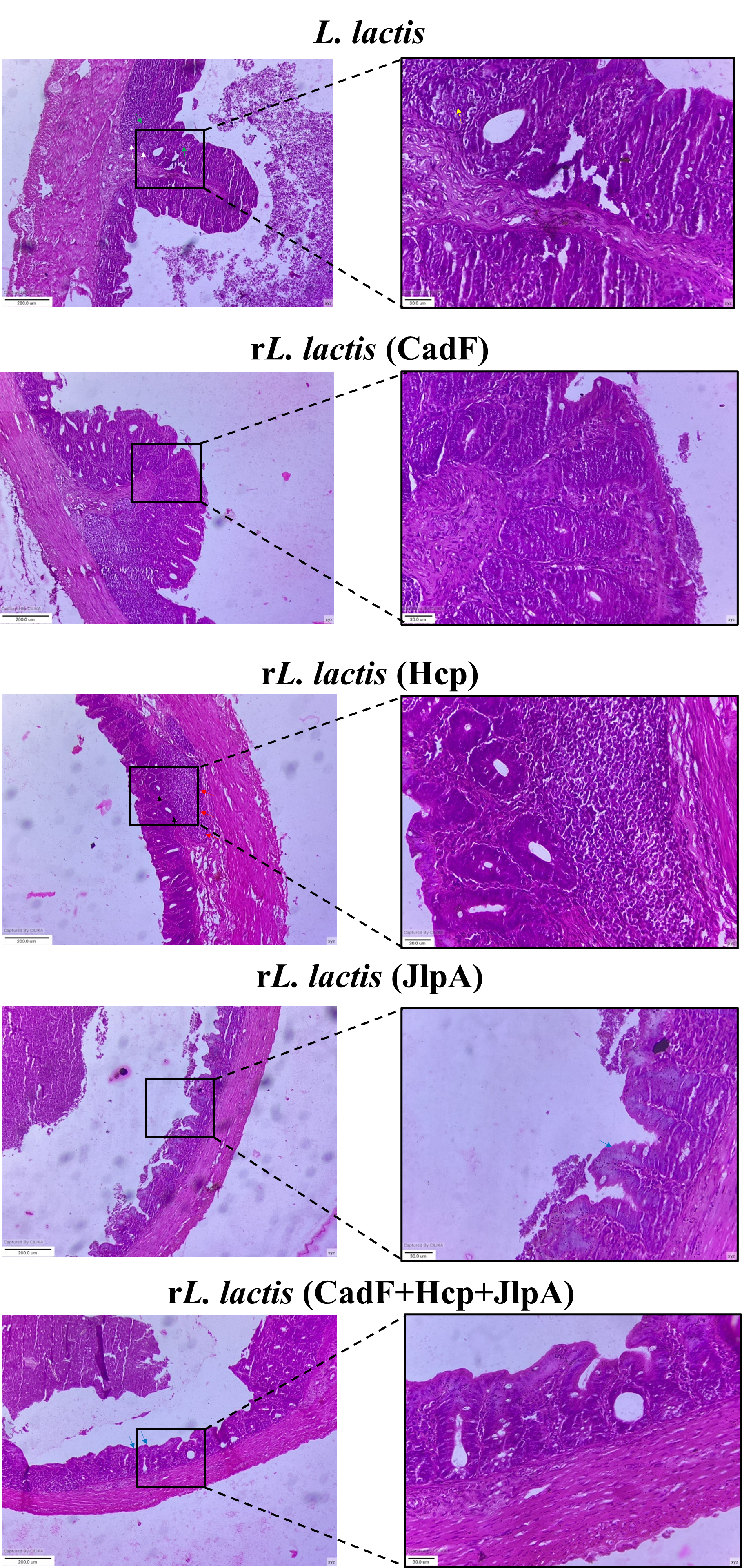

**Figure S5:** **Additional images of tissue sections showing histopathological changes** of cecal tissue collected from birds at day 7 post-infection with *C. jejuni* (TGH 9011). Green arrow: cellular infiltration, yellow arrow- crypt disruption, white arrow- tissue necrosis, red arrow- lymphoid accumulation, black arrow- well-organised crypt, blue arrow- intact epithelia.

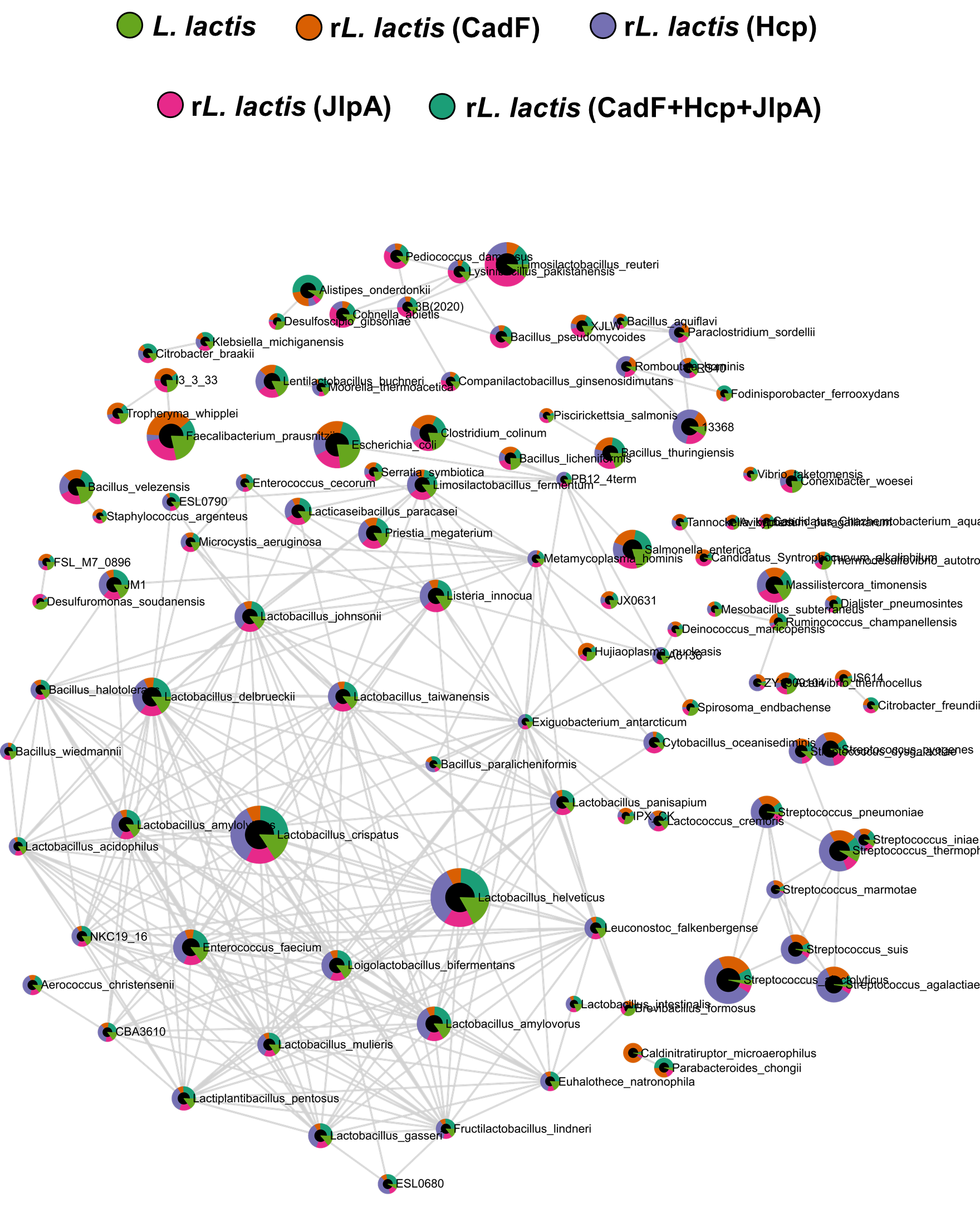

**Figure S6:** **Correlation network analysis of cecal microbiota.** The correlation network was analysed using the Sparse Correlations for Compositional Data (SparCC) approach. The microbial correlation network illustrated intricate patterns of interaction, where nodes are taxa and edges are notable correlations. Each node has a pie chart displaying the relative occurrence of each group in shared taxa.

| **WT *L. lactis* vs r*L. lactis* (CadF)** | | | | | | | | | | | | | |
| --- | --- | --- | --- | --- | --- | --- | --- | --- | --- | --- | --- | --- | --- |
| **Phylum** | **Log2FC** | **St. Error** | | | | **P-value** | | | | **FDR** | | | |
| Planctomycetota | -1.67 | 0.297 | | | | 4.97E-05 | | | | 0.00172 | | | |
| Candidatus_Saccharibacteria | -1.35 | 0.391 | | | | 0.00359 | | | | 0.0747 | | | |
| Cyanobacteriota | 0.397 | 0.186 | | | | 0.0493 | | | | 0.265 | | | |
| Fusobacteriota | 0.626 | 0.49 | | | | 0.221 | | | | 0.588 | | | |
| Bacteroidota | -0.78 | 0.839 | | | | 0.368 | | | | 0.693 | | | |
| Chlamydiota | 0.647 | 0.699 | | | | 0.369 | | | | 0.693 | | | |
| Chordata | 0.237 | 0.273 | | | | 0.4 | | | | 0.716 | | | |
| Bacillota | 0.0129 | 0.0188 | | | | 0.503 | | | | 0.818 | | | |
| Actinomycetota | 0.121 | 0.196 | | | | 0.546 | | | | 0.847 | | | |
| Candidatus_Bipolaricaulota | -0.376 | 0.605 | | | | 0.544 | | | | 0.847 | | | |
| Nitrospirota | 0.617 | 0.963 | | | | 0.531 | | | | 0.847 | | | |
| Aquificota | -0.149 | 0.3 | | | | 0.626 | | | | 0.88 | | | |
| Spirochaetota | 0.141 | 0.264 | | | | 0.602 | | | | 0.88 | | | |
| Synergistota | -0.355 | 0.684 | | | | 0.612 | | | | 0.88 | | | |
| Campylobacterota | -0.182 | 0.42 | | | | 0.671 | | | | 0.907 | | | |
| Candidatus_Omnitrophota | -0.11 | 0.375 | | | | 0.774 | | | | 0.938 | | | |
| Ignavibacteriota | -0.219 | 0.795 | | | | 0.787 | | | | 0.938 | | | |
| Pseudomonadota | 0.0946 | 0.313 | | | | 0.766 | | | | 0.938 | | | |
| Thermodesulfobacteriota | 0.259 | 0.905 | | | | 0.779 | | | | 0.938 | | | |
| Unknown | -0.0494 | 0.186 | | | | 0.794 | | | | 0.938 | | | |
| Acidobacteriota | 0.187 | 0.859 | | | | 0.831 | | | | 0.967 | | | |
| Bacteria_none_none | 0.135 | 0.907 | | | | 0.883 | | | | 0.967 | | | |
| Bdellovibrionota | 0.0905 | 0.586 | | | | 0.879 | | | | 0.967 | | | |
| Mycoplasmatota | -0.0416 | 0.404 | | | | 0.919 | | | | 0.976 | | | |
| Deinococcota | 0.0156 | 0.68 | | | | 0.982 | | | | 0.994 | | | |
| Thermotogota | -0.00675 | 0.844 | | | | 0.994 | | | | 0.994 | | | |
| **WT *L. lactis* vs r*L. lactis* (Hcp)** | | | | | | | | | | | | | |
| **Phylum** | **Log_2_FC** | | **St. Error** | | | | **P-value** | | | | **FDR** | | |
| Planctomycetota | -2.31 | | 0.297 | | | | 1.22E-06 | | | | 4.24E-05 | | |
| Candidatus_Saccharibacteria | -1.47 | | 0.391 | | | | 0.00187 | | | | 0.0389 | | |
| Actinomycetota | 0.622 | | 0.196 | | | | 0.00627 | | | | 0.0712 | | |
| Candidatus_Omnitrophota | 1.09 | | 0.375 | | | | 0.011 | | | | 0.0712 | | |
| Chlamydiota | 2.22 | | 0.699 | | | | 0.00624 | | | | 0.0712 | | |
| Chordata | 0.796 | | 0.273 | | | | 0.0107 | | | | 0.0712 | | |
| Cyanobacteriota | 0.556 | | 0.186 | | | | 0.00909 | | | | 0.0712 | | |
| Aquificota | 0.839 | | 0.3 | | | | 0.0135 | | | | 0.078 | | |
| Campylobacterota | 1.02 | | 0.42 | | | | 0.0276 | | | | 0.125 | | |
| Thermodesulfobacteriota | 2.22 | | 0.905 | | | | 0.027 | | | | 0.125 | | |
| Pseudomonadota | 0.749 | | 0.313 | | | | 0.0302 | | | | 0.129 | | |
| Bacillota | -0.0435 | | 0.0188 | | | | 0.0358 | | | | 0.138 | | |
| Nitrospirota | 2.09 | | 0.963 | | | | 0.0465 | | | | 0.158 | | |
| Ignavibacteriota | 1.62 | | 0.795 | | | | 0.0599 | | | | 0.164 | | |
| Unknown | 0.393 | | 0.186 | | | | 0.0517 | | | | 0.164 | | |
| Mycoplasmatota | 0.748 | | 0.404 | | | | 0.0838 | | | | 0.203 | | |
| Bdellovibrionota | 1.02 | | 0.586 | | | | 0.101 | | | | 0.224 | | |
| Bacteria_none_none | 1.54 | | 0.907 | | | | 0.11 | | | | 0.233 | | |
| Fusobacteriota | 0.827 | | 0.49 | | | | 0.112 | | | | 0.233 | | |
| Acidobacteriota | 0.98 | | 0.859 | | | | 0.272 | | | | 0.404 | | |
| Candidatus_Bipolaricaulota | 0.492 | | 0.605 | | | | 0.429 | | | | 0.56 | | |
| Synergistota | -0.331 | | 0.684 | | | | 0.635 | | | | 0.76 | | |
| Thermotogota | 0.367 | | 0.844 | | | | 0.67 | | | | 0.774 | | |
| Deinococcota | 0.155 | | 0.68 | | | | 0.822 | | | | 0.891 | | |
| Bacteroidota | -0.0994 | | 0.839 | | | | 0.907 | | | | 0.933 | | |
| Spirochaetota | 0.0086 | | 0.264 | | | | 0.974 | | | | 0.974 | | |
| **WT *L. lactis* vs r*L. lactis* (JlpA)** | | | | | | | | | | | | | |
| **Phylum** | **Log2FC** | | | | **St. Error** | | | | **P-value** | | | **FDR** | |
| Chlamydiota | 1.46 | | | | 0.699 | | | | 0.0539 | | | 0.538 | |
| Chordata | 0.511 | | | | 0.273 | | | | 0.081 | | | 0.579 | |
| Cyanobacteriota | 0.334 | | | | 0.186 | | | | 0.0927 | | | 0.579 | |
| Deinococcota | -0.699 | | | | 0.68 | | | | 0.32 | | | 0.649 | |
| Fusobacteriota | 0.721 | | | | 0.49 | | | | 0.162 | | | 0.649 | |
| Ignavibacteriota | 0.986 | | | | 0.795 | | | | 0.234 | | | 0.649 | |
| Nitrospirota | 1.29 | | | | 0.963 | | | | 0.199 | | | 0.649 | |
| Synergistota | 0.783 | | | | 0.684 | | | | 0.271 | | | 0.649 | |
| Thermodesulfobacteriota | 0.973 | | | | 0.905 | | | | 0.299 | | | 0.649 | |
| Thermotogota | -0.812 | | | | 0.844 | | | | 0.351 | | | 0.666 | |
| Candidatus_Omnitrophota | 0.314 | | | | 0.375 | | | | 0.415 | | | 0.73 | |
| Unknown | 0.149 | | | | 0.186 | | | | 0.436 | | | 0.73 | |
| Aquificota | 0.219 | | | | 0.3 | | | | 0.477 | | | 0.763 | |
| Campylobacterota | 0.257 | | | | 0.42 | | | | 0.55 | | | 0.823 | |
| Bacteroidota | 0.383 | | | | 0.839 | | | | 0.655 | | | 0.884 | |
| Spirochaetota | 0.132 | | | | 0.264 | | | | 0.624 | | | 0.884 | |
| Bacillota | -0.0077 | | | | 0.0188 | | | | 0.688 | | | 0.896 | |
| Pseudomonadota | 0.128 | | | | 0.313 | | | | 0.689 | | | 0.896 | |
| Bdellovibrionota | 0.195 | | | | 0.586 | | | | 0.744 | | | 0.922 | |
| Acidobacteriota | 0.167 | | | | 0.859 | | | | 0.849 | | | 0.98 | |
| Candidatus_Bipolaricaulota | 0.12 | | | | 0.605 | | | | 0.846 | | | 0.98 | |
| Actinomycetota | 0.0273 | | | | 0.196 | | | | 0.891 | | | 0.986 | |
| Bacteria_none_none | 0.128 | | | | 0.907 | | | | 0.889 | | | 0.986 | |
| Candidatus_Saccharibacteria | 6.58E-15 | | | | 0.391 | | | | 1 | | | 1 | |
| Mycoplasmatota | 0.0274 | | | | 0.404 | | | | 0.947 | | | 1 | |
| Planctomycetota | 8.69E-15 | | | | 0.297 | | | | 1 | | | 1 | |
| **WT *L. lactis* vs r*L. lactis* (CadF+Hcp+JlpA)** | | | | | | | | | | | | | |
| **Phylum** | **Log2FC** | | | **St. Error** | | | | **P-value** | | | | | **FDR** |
| Fusobacteriota | 1.83 | | | 0.49 | | | | 0.00199 | | | | | 0.069 |
| Synergistota | 1.66 | | | 0.684 | | | | 0.0285 | | | | | 0.269 |
| Actinomycetota | 0.449 | | | 0.196 | | | | 0.037 | | | | | 0.278 |
| Thermodesulfobacteriota | 2.12 | | | 0.905 | | | | 0.0334 | | | | | 0.278 |
| Chlamydiota | 1.46 | | | 0.699 | | | | 0.0535 | | | | | 0.304 |
| Acidobacteriota | 1.65 | | | 0.859 | | | | 0.0744 | | | | | 0.352 |
| Bacteroidota | -1.5 | | | 0.839 | | | | 0.0936 | | | | | 0.363 |
| Spirochaetota | 0.47 | | | 0.264 | | | | 0.095 | | | | | 0.363 |
| Unknown | 0.336 | | | 0.186 | | | | 0.091 | | | | | 0.363 |
| Bdellovibrionota | 0.935 | | | 0.586 | | | | 0.131 | | | | | 0.371 |
| Chordata | 0.393 | | | 0.273 | | | | 0.171 | | | | | 0.42 |
| Aquificota | 0.366 | | | 0.3 | | | | 0.241 | | | | | 0.494 |
| Campylobacterota | 0.512 | | | 0.42 | | | | 0.242 | | | | | 0.494 |
| Bacteria_none_none | 0.968 | | | 0.907 | | | | 0.303 | | | | | 0.533 |
| Nitrospirota | 0.947 | | | 0.963 | | | | 0.341 | | | | | 0.581 |
| Candidatus_Bipolaricaulota | 0.586 | | | 0.605 | | | | 0.348 | | | | | 0.584 |
| Candidatus_Omnitrophota | 0.342 | | | 0.375 | | | | 0.377 | | | | | 0.603 |
| Ignavibacteriota | 0.699 | | | 0.795 | | | | 0.393 | | | | | 0.61 |
| Pseudomonadota | 0.217 | | | 0.313 | | | | 0.499 | | | | | 0.73 |
| Candidatus_Saccharibacteria | -0.25 | | | 0.391 | | | | 0.532 | | | | | 0.754 |
| Deinococcota | 0.359 | | | 0.68 | | | | 0.605 | | | | | 0.798 |
| Cyanobacteriota | 0.0682 | | | 0.186 | | | | 0.719 | | | | | 0.865 |
| Mycoplasmatota | 0.104 | | | 0.404 | | | | 0.8 | | | | | 0.895 |
| Bacillota | 0.003 | | | 0.0188 | | | | 0.876 | | | | | 0.945 |
| Thermotogota | 0.101 | | | 0.844 | | | | 0.907 | | | | | 0.952 |
| Planctomycetota | 8.69E-15 | | | 0.297 | | | | 1 | | | | | 1 |

**Table S5: Multiple linear regression with covariate adjustment of cecal microbiota at the phylum level.** The MicrobiomeAnalyst (2.0), freely available software, was used to calculate the multiple regression between experimental groups. The tool uses general linear models to find associations between microbial features and experimental metadata using MaAsLin2. A linear model was fitted to each microbial feature, including the primary metadata and covariates.

| **Chao1** | | | |
| --- | --- | --- | --- |
| **Pair** | **Statistic** | **P-value** | **FDR** |
| L._lactis vs rL._lactis_(CadF) | 0.14163 | 0.89218 | 0.89218 |
| L._lactis vs rL._lactis_(Hcp) | 0.8236 | 0.44196 | 0.89218 |
| L._lactis vs rL._lactis_(JlpA) | 0.58508 | 0.57995 | 0.89218 |
| L._lactis vs rL._lactis_(CadF+Hcp+JlpA) | 0.80197 | 0.46031 | 0.89218 |
| rL._lactis_(CadF) vs rL._lactis_(Hcp) | 0.61099 | 0.56382 | 0.89218 |
| rL._lactis_(CadF) vs rL._lactis_(JlpA) | 0.37515 | 0.72137 | 0.89218 |
| rL._lactis_(CadF) vs rL._lactis_(CadF+Hcp+JlpA) | 0.65181 | 0.54128 | 0.89218 |
| rL._lactis_(Hcp) vs rL._lactis_(JlpA) | -0.29992 | 0.77467 | 0.89218 |
| rL._lactis_(Hcp) vs rL._lactis_(CadF+Hcp+JlpA) | 0.18158 | 0.8629 | 0.89218 |
| rL._lactis_(JlpA) vs rL._lactis_(CadF+Hcp+JlpA) | 0.41075 | 0.69974 | 0.89218 |
| **Shannon** | | | |
| **Pair** | **Statistic** | **P-value** | **FDR** |
| L._lactis vs rL._lactis_(CadF) | -0.63739 | 0.55477 | 0.69347 |
| L._lactis vs rL._lactis_(Hcp) | 3.1998 | 0.041542 | 0.20771 |
| L._lactis vs rL._lactis_(JlpA) | 0.26804 | 0.80488 | 0.80488 |
| L._lactis vs L._lactis_(CadF+Hcp+JlpA) | 1.6584 | 0.18948 | 0.37896 |
| rL._lactis_(CadF) vs rL._lactis_(Hcp) | 3.281 | 0.026725 | 0.20771 |
| rL._lactis_(CadF) vs rL._lactis_(JlpA) | 0.50941 | 0.63835 | 0.70927 |
| rL._lactis_(CadF) vsrL._lactis_(CadF+Hcp+JlpA] | 1.8335 | 0.1442 | 0.3605 |
| rL._lactis_(Hcp) vs rL._lactis_(JlpA) | -1.7943 | 0.1262 | 0.3605 |
| rL._lactis_(Hcp) vs rL._lactis_(CadF+Hcp+JlpA) | -0.64014 | 0.54746 | 0.69347 |
| rL._lactis_(JlpA) vs rL._lactis_(CadF+Hop+JlpA) | 1.0134 | 0.35001 | 0.58335 |
| **Simpson** | | | |
| **Pair** | **Statistic** | **P-value** | **FDR** |
| L._lactis vs rL._lactis_(CadF) | -0.47138 | 0.65435 | 0.65435 |
| L._lactis rL._lactis_(Hcp) | 2.7161 | 0.065585 | 0.32792 |
| L._lactis vs rL._lactis_(JlpA) | 0.64174 | 0.564 | 0.64538 |
| L._lactis vs rL._lactis_(CadF+Hcp+JlpA) | 1.6283 | 0.19547 | 0.48868 |
| rL._lactis_(CadF) vs rL._lactis_(Hcp) | 2.8507 | 0.056157 | 0.32792 |
| rL._lactis_(CadF) vs rL._lactis_(JlpA) | 0.76849 | 0.49385 | 0.64538 |
| rL._lactis_(CadF) vs L._lactis_(CadF+Hcp+JlpA) | 1.7559 | 0.16882 | 0.48868 |
| rL._lactis_(Hcp) vs rL._lactis_(JlpA) | -1.2704 | 0.25258 | 0.50515 |
| rL._lactis_(Hcp) vs rL._lactis_(CadF+Hcp+JlpA) | -0.58403 | 0.58084 | 0.64538 |
| rL._lactis_(JlpA) vs rL._lactis_(CadF+Hcp+JlpA) | 0.66511 | 0.53078 | 0.64538 |
| **Fisher** | | | |
| **Pair** | **Statistic** | **P-value** | **FDR** |
| L._lactis vs rL._lactis_(CadF) | 0.21766 | 0.83569 | 0.92647 |
| L._lactis vs rL._lactis_(Hcp) | 1.2201 | 0.27254 | 0.87595 |
| L._lactis vs rL._lactis_(JlpA) | 0.75183 | 0.48067 | 0.87595 |
| L._lactis vs rL._lactis_(CadF+Hcp+JlpA) | 0.9187 | 0.40968 | 0.87595 |
| rL._lactis_(CadF) vs rL._lactis_(Hcp) | 0.8621 | 0.42176 | 0.87595 |
| rL._lactis_(CadF) vs rL._lactis_(JlpA) | 0.40484 | 0.70076 | 0.87595 |
| rL._lactis_(CadF) vs L._lactis_(CadF+Hcp+JlpA) | 0.71818 | 0.50531 | 0.87595 |
| rL._lactis_(Hcp) vs rL._lactis_(JlpA) | -0.5711 | 0.59011 | 0.87595 |
| rL._lactis_(Hcp) vs rL._lactis_(CadF+Hcp+JlpA) | 0.097173 | 0.92647 | 0.92647 |
| rL._lactis_(JlpA) vs rL._lactis_(CadF+Hcp+JlpA) | 0.48823 | 0.65006 | 0.87595 |

**Table S6:** The table summarises the post-hoc pairwise comparison (multi-group) of alpha diversity indexes (Chao1, Shannon, Simpson and Fisher). Regular Welch t-tests/ANOVA were performed for each pair. The multi-testing adjustment is based on the Benjamini-Hochberg procedure (FDR).

| **Beta diversity (PERMANOVA)** | | | | |
| --- | --- | --- | --- | --- |
| **Pair** | **F-value** | **R-squared** | **P-value** | **FDR** |
| L._lactis vs rL._lactis_(CadF) | 1.2114 | 0.16798 | 0.337 | 0.42125 |
| L._lactis vs rL._lactis_(Hcp) | 7.2456 | 0.54702 | 0.031 | 0.155 |
| L._lactis vs rL._lactis_(JlpA) | 0.7802 | 0.11507 | 0.614 | 0.614 |
| L._lactis vs rL._lactis_(CadF+Hcp+JlpA) | 1.4239 | 0.1918 | 0.271 | 0.38714 |
| rL._lactis_(CadF) vs rL._lactis_(Hcp) | 5.9128 | 0.49634 | 0.075 | 0.156 |
| rL._lactis_(CadF) vs rL._lactis_(JlpA) | 0.73606 | 0.10927 | 0.607 | 0.614 |
| rL._lactis_(CadF) vs rL._lactis_(CadF+Hcp+JlpA) | 2.7123 | 0.31132 | 0.057 | 0.156 |
| rL._lactis_(Hcp) vs rL._lactis_(JlpA) | 4.2234 | 0.41311 | 0.078 | 0.156 |
| rL._lactis_(Hcp) vs rL._lactis_(CadF+Hcp+JlpA) | 3.3224 | 0.35639 | 0.023 | 0.155 |
| rL._lactis_(JlpA) vs rL._lactis_(CadF+Hcp+JlpA) | 1.6073 | 0.21129 | 0.186 | 0.31 |

**Table S7:** The table summarises the results of the pairwise PERMANOVA analysis of beta diversity. The multi-testing adjustment is based on the Benjamini-Hochberg procedure (FDR).
